# When Multi-Voxel Pattern Similarity and Global Activation are Intertwined: Approaches to Disentangling Correlation from Activation

**DOI:** 10.1101/2023.05.29.542175

**Authors:** Karen F. LaRocque, Tyler H. Davis, Jeanette A. Mumford, Russell A. Poldrack, Anthony D. Wagner

**Affiliations:** Department of Psychology, Stanford University; Wu Tsai Neurosciences Institute, Stanford University; Independent Scientist, California

## Abstract

Pattern similarity analysis, which uses correlation to examine similarities between neural activation patterns evoked by different trials or conditions, is often leveraged to test hypotheses not easily answerable with univariate comparisons, such as *how* events are represented or processed and the *relationships* between representations or processing of events. In principle, univariate analyses of global activation and multivariate analyses of pattern similarity can be used to answer substantively different questions about psychological and neural processing. For this to hold, it is necessary that pattern similarity estimates are not contaminated by differences in global activation across experimental events. Here, we report simulated data that demonstrate that global activation and pattern similarity (as assessed by correlation), although theoretically independent, are often intertwined. We present two plausible scenarios that illustrate how condition-specific changes in global activation can elicit condition-specific increases in pattern similarity by interacting with underlying across-voxel activation patterns. First, we consider a scenario in which a target region contains subpopulations of voxels such that only some voxels in a region are sensitive to a psychological variable and the remaining voxels are not modulated by this variable. In this scenario, this spatial pattern of responsive and unresponsive voxels *adds new, shared across-voxel variability* for events in the ‘active’ condition, thereby increasing pattern similarity between these events. Second, we consider a scenario in which trials from all conditions elicit a shared across-voxel pattern of activation, but this shared across-voxel pattern is amplified for trials within one condition due to greater global activation. In this scenario, the change in activation for a given condition *increases the ability to detect pre-existing, shared across-voxel variability* across events in that condition, thereby increasing pattern similarity between these events. Given the observed influence of global activation on pattern similarity, we then assess whether it is possible to statistically separate the contributions of global activation and pattern similarity to observed activation patterns (using regression approaches, matching activation across conditions, and inclusion of control conditions). Additional simulations demonstrate that use of these techniques is not always effective in removing the influence of global activation on pattern similarity ––the efficacy of these techniques depends on a variety of signal parameters that will likely vary across experiments and participants, highlighting the need for tailored control analyses that are targeted at addressing the particular hypotheses and potential global activation confounds of a given experiment.

[**Note:** The reported simulations and this resulting white paper were generated in 2014. We share, without update, this paper given the continued relevance of understanding and controlling for global activation confounds when conducting multi-variate pattern analyses.]

## Introduction

Application of multi-voxel pattern analysis (MVPA) has greatly expanded the types of questions that can be addressed with functional magnetic resonance imaging (fMRI). In addition to enabling detection of neural signals operating at a higher spatial frequency than is detectable by traditional univariate analyses (e.g., Haxby et al., 2001; Haynes and Rees, 2006; Jimura and Poldrack, 2012; Kamitani and Tong, 2005; Norman et al., 2006), MVPA allows for fine-grained tests of the relationships between activation patterns elicited by distinct events (e.g., Edelman et al., 1998; Kriegeskorte et al., 2008). As such, MVPA provides a level of analysis that compliments traditional univariate tests of global signal change (for review see Coutanche, 2013; Davis and Poldrack, 2013; Kriegeskorte and Kievit, 2013). Particularly useful here is pattern similarity analysis, which examines similarities between activation patterns evoked by different trials or conditions within a task (e.g., Edelman et al., 1998; Kriegeskorte et al., 2008). In these applications, the goal is often to test hypotheses not easily answerable with univariate comparisons of global increases or decreases in activation within a region, such as *how* events are represented or processed and the *relationships* between representations or processing of events (c.f., fMRI-adaptation; Grill-Spector and Malach, 2001).

By way of example, consider a hypothetical region involved in processing a psychological stimulus dimension, such as scariness. Increased attention to scariness may increase processing or engagement in this region, resulting in a global increase in activation within the region. However, increased attention to scariness may also evoke independent changes in the region’s similarity relationships between patterns of activation elicited by stimuli that vary along the dimension of scariness. For example, attention to scariness may make activation patterns elicited by stimuli of similar scariness more similar, whereas the opposite may occur for stimuli that differ in their degree of scariness. In this example, there would be two signals underlying activation within a region, with global changes in activation indicating overall engagement of attention to scariness, and changes in pattern similarity indicating how attention to scariness has changed the weighting of information representation. The recent literature contains a number of examples where these two types of signals may exist within a single task, often in contexts testing cognitive theories of representation using BOLD fMRI data (e.g., Davis and Poldrack, 2013; Kriegeskorte and Kievit, 2013).

In principle, univariate analyses of global activation and multivariate analyses of pattern similarity can be used to answer substantively different questions about psychological and neural processing. For this to hold, however, it is necessary that pattern similarity estimates are not contaminated by differences in global activation across experimental events. Indeed, if global activation tracks a theoretically separate component of the neural signal from that revealed by pattern similarity analysis, it is important to remove this component so that it does not interfere with measurements of pattern similarity. In our example, a large change in activation within a region due to shifts in attention to scariness could mask smaller changes in the weighting of information within the region. The use of correlation as a similarity measure has been argued to alleviate this problem as it removes information about mean activation across voxels for a given event, thus theoretically enabling independent assessment of similarity relationships across events. However, this critical assumption requires formal evaluation, as initial observations suggest it may not uniformly hold (e.g., LaRocque et al., 2013).

Here, we use simulated data to demonstrate that global activation and pattern similarity (as assessed by correlation), although theoretically independent, are often intertwined inpractice. We start with the foundation that correlations are measures of the degree of shared across-voxel variability in the activation patterns elicited across events (items or conditions) in a task (Davis et al., 2014). Critically, any factor that either (a) increases shared across-voxel variability or (b) increases the ability to detect pre-existing shared across-voxel variability between activation patterns across two events will increase correlation-based measures of the pattern similarity of those two events.

We present two plausible scenarios that illustrate how condition-specific changes in global activation can elicit condition-specific increases in pattern similarity by interacting with underlying across-voxel activation patterns. First, we consider a scenario in which a target region contains subpopulations of voxels such that only some voxels in a region are sensitive to a psychological variable and the remaining voxels are not modulated by this variable. For example, a region may span multiple functional subpopulations of voxels (Friston et al., 2006; Grill-Spector & Malach, 2001), or may contain subpopulations of voxels that are unresponsive to a task due to susceptibility artifacts or partial voluming effects. In this scenario, this spatial pattern of responsive and unresponsive voxels *adds new, shared across-voxel variability* for events in the ‘active’ condition, thereby increasing pattern similarity between these events. Second, we consider a scenario in which trials from all conditions elicit a shared across-voxel pattern of activation, but this shared across-voxel pattern is amplified for trials within one condition due to greater global activation. In other words, global signal serves to increase the gain on the expression of an activation pattern by multiplying the baseline activation in each voxel by a scalar. Such a scaling of responses across conditions has been observed in contexts comparing novel and repeated stimuli (e.g., Grill-Spector et al., 2006; Li et al., 1993; Weiner et al., 2010) or attended and unattended stimuli. In this scenario, the change in activation for a given condition *increases the ability to detect pre-existing, shared across-voxel variability* across events in that condition, thereby increasing pattern similarity between these events.

Given the observed influence of global activation on pattern similarity, we then assess whether it is possible to statistically separate the contributions of global activation and pattern similarity to observed activation patterns. Here, we use several traditional statistical techniques — regression approaches, matching activation across conditions, and inclusion of control conditions. Through additional simulations, we demonstrate that use of these techniques is not always effective in removing the influence of global activation on pattern similarity. Specifically, we demonstrate that the efficacy of these techniques depends on a variety of signal parameters that will likely vary across experiments and participants, highlighting the need for tailored control analyses that are targeted at addressing the particular hypotheses and potential global activation confounds of a given experiment.

## Methods

### General simulation approach

To examine the relationship between global activation and pattern similarity, we considered the ‘attention to scariness’ scenario outlined in the Introduction, in which a region is hypothesized to process the psychological dimension of ‘scariness’. The region may show an increase in global activation (reflecting differential engagement) when processing scary relative to non-scary stimuli, but also may independently perform a transformation on its inputs such that the degree of scariness of a stimulus is represented in the multidimensional space of across-voxel pattern of activation elicited by the stimulus. Critically, in the current simulations, the region’s signal is generated to exhibit only an effect of scariness on global levels of activation, such that (a) activation is higher for scary relative to non-scary stimuli, and (b) across-voxel pattern similarity does not code for scariness beyond these changes in global activation. We then ask whether the observed pattern of data reflects this ‘ground truth’, i.e., whether the data demonstrate an effect of condition on activation but not on pattern similarity. Note that in the full scenario in the Introduction, these results would be compared to a ‘no attention to scariness’ condition in which neither an effect of scariness on activation or on pattern similarity is observed.

All simulations were performed using R (R core team, 2014). Data were simulated as the sum of a ground-truth pattern of activation across voxels for each stimulus, global activation increases due to stimulus condition (scary/non-scary), and normally distributed noise (**Figure 1**). Ground-truth data were simulated as activation values for 50 stimuli across 200 voxels. A ground-truth across-voxel pattern of activation for each stimulus was drawn from a multivariate normal distribution. Unless otherwise stated, the distribution was such that the mean activation for each voxel was 1 (all units arbitrary), the across-voxel variance for each stimulus was 1, and the covariance across stimuli was .50. Note that because the mean across-voxel variance was 1, the ground-truth covariance is equivalent to the ground-truth correlation in all simulations. Half of the stimuli were assigned to the ‘non-active’ condition (here, the ‘non-scary’ condition) and half were assigned to the ‘active’ condition (the ‘scary condition). Condition-specific activation increases were assigned to the active condition in one of three ways: in the *uniform* case, every voxel received an activation increase that was constant across all stimuli; in the *subset* case, 50% of the voxels received an activation increase that was constant across all stimuli; in the *amplified* case, every voxel received a stimulus-specific activation increase equal to a multiple of the ground-truth activation value of that voxel for that stimulus (because the mean activation was greater than zero this produced an overall increase in activation). The amount of activation (or the multiple of the amount of activation) was parametrically varied from 0 to 3 in steps of 0.5. Finally, unless otherwise stated, a noise value drawn from a normal distribution with a mean of 0 and a variance of 1 was added to each voxel separately for each stimulus.

**Figure 1.**
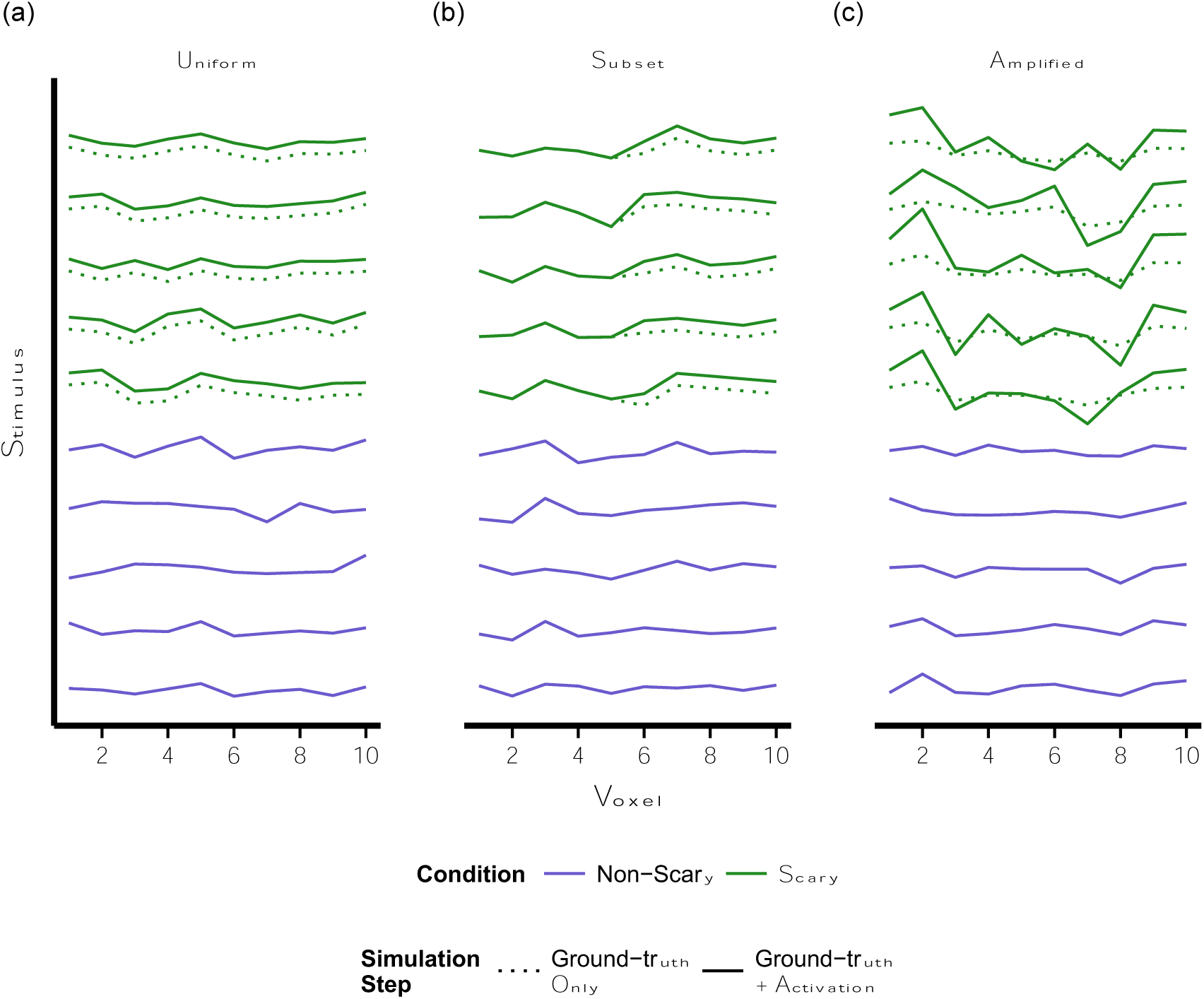
Simulation approach. Activation was added to scary stimuli as **(a)** a uniform increase across all voxels (*uniform* case); this increase in activation has no effect on shared across-voxel variability across conditions, **(b)** a uniform increase for 50% of the voxels (*subset* case); this increase in activation adds new shared across-voxel variability to the scary stimuli, or **(c)** a stimulus-specific increase across all voxels that is equal to a multiple of the baseline response to each stimulus (*amplified* case); this increase in activation amplifies and allows for detection of the pre-existing shared across-voxel variability for scary stimuli (in the face of noise). Note that randomly distributed noise was also added to the final simulated patterns of response (not pictured here).

In all cases, our measure of interest was whether pattern similarity differed across pairs of stimuli drawn both from the non-active (non-scary) condition (NN pairs), both from the active (scary) condition (SS pairs), or one stimulus drawn from the active condition and the other from the non-active condition (SN pairs). To test this, we computed correlations of the across-voxel pattern of response between all stimulus pairs and took the mean of these correlations for NN pairs, SS pairs, and SN pairs separately.

This process was repeated 100 times for each simulation; all plots show mean correlation values and standard deviations across these 100 simulations.

### Simulations with distributed activation increases across stimuli

In several simulations (those reported in **Figures 4, S2, S3, S4, & S5**) we included additional variability in the activation increases applied to stimuli within both conditions. To accomplish this, we allowed the activation applied to each stimulus in the active condition to vary such that it was distributed according to a χ^2^ distribution with a mean that parametrically varied from 0 to 3 in steps of 0.5. The use of a χ^2^ distribution ensured that activation increased for all stimuli in the active condition and that the variance of the amount of activation added across stimuli increased as the mean amount of activation increased.

### Simulations involving a control condition

A final set of simulations mimicked a design in which single stimuli within the two conditions (ten stimuli; five per condition) are presented multiple times (five times each). The mean covariance across stimuli was allowed to vary independently of the mean covariance between repetitions of the same stimulus. The mean covariance between repetitions of the same stimulus (within-stimulus covariance) remained set at .50, while the mean covariance between distinct stimuli (across-stimulus covariance) was set at either .25 or .50. In these simulations, pattern similarity was computed separately for pairs of repeated (within-stimulus) versus distinct (across-stimulus) stimuli separately for the non-active and active conditions.

### Procedures correcting for the influence of activation on correlation

We applied several statistical adjustments to the data generated from the simulation procedures described above. Each adjustment process was repeated 100 times; all plots show mean adjusted values and standard deviations across these 100 simulations.

First, we examined whether effects of activation could be removed after correlations had already been computed. For each pair of stimuli, we calculated (a) the correlation between those two stimuli and (b) the sum of the mean activation for each stimulus across all voxels. We regressed correlations on activation irrespective of pair type (e.g., whether each correlation was a NN, SS, or SN pair). We then binned the residuals by pair type and examined whether the means of these residuals differed by pair type.

Second, we examined whether effects of activation could be removed from individual voxels before correlations are computed. We mean centered each voxel by computing its mean activation across the entire set of stimuli and subtracted this mean from its response to each individual stimulus. We then used the remaining responses for each voxel to compute correlations between each pair of stimuli and examined whether the mean correlation differed by pair type.

Third, we examined whether condition-specific differences in correlations could be attenuated if stimuli from the two conditions were matched on activation via subsampling. In a first set of simulations we removed voxels that showed a greater response to stimuli in one condition relative to the other. Specifically, we performed a t-test on each voxel comparing activation for stimuli from the non-active condition to activation for stimuli from the active condition. If the p-value yielded by this test was less than a set statistical threshold (set to .05 or .50), then the voxel was removed. If five or more voxels remained following this procedure, pair-wise correlations were then performed across the remaining voxels and mean correlations were computed for each pair type; if fewer than five voxels remained, the simulation was ended and not included in the results. In a second set of simulations we subsampled stimuli that were matched on mean level of activation across all voxels. Specifically, we used the R package ‘MatchIt’ (discarding units from both conditions that fell outside of the common support and then using nearest neighbor matching; Ho et al., 2011; 2007) to identify and retain a subset of stimuli that were matched on activation. If 10 or more stimuli remained in each condition following this procedure, pair-wise correlations were then performed across the remaining stimuli and mean correlations were computed for each pair type; if fewer then 10 stimuli remained in either condition, the simulation was ended and not included in the results.

## Results

### Condition-specific increases in global activation can alter multivariate pattern similarity

First, we examined whether condition-specific increases in global activation can produce condition-specific changes in across-voxel pattern similarity. In other words, we examined whether what would traditionally be labeled an effect of condition on activation could produce what appears to be an effect of condition on pattern similarity. To do so, we first simulated ground-truth patterns of stimulus-specific voxel responses to scary and non-scary stimuli. These across-voxel patterns were either uncorrelated (correlation = 0.00) or correlated (correlation = 0.50). Importantly, these ground-truth correlations between stimuli did not differ as a function of stimulus condition. We then simulated, on top of these patterns, either a global increase in activation (scary stimuli) or no global change in activation (non-scary stimuli). The global increase in activation for the scary stimuli took one of three forms: a uniform increase across all voxels (*uniform*), a uniform increase in only a subset of voxels (*subset*), or a scaled increase across all voxels that amplified pre-existing patterns of responses across voxels (*amplified*). The final observed pattern across voxels for each stimulus was the sum of the ground-truth pattern of response across voxels, any effects of activation, and normally distributed noise (**Figure 1**). We then assessed whether the mean correlation of the observed across-voxel patterns of response for two stimuli, our measure of pattern similarity, differed for pairs of stimuli drawn from the same condition (scary—scary (SS) or non-scary—non-scary (NN)) relative to pairs of stimuli drawn from different conditions (scary—non-scary (SN)). Importantly, any increased pattern similarity for within-condition pairs of stimuli (SS and / or NN) relative to across-condition pairs of stimuli (SN) would be mistaken as evidence that the region is coding for scariness in a manner that extends beyond a change in global activation, even though the increased similarity would be directly due to the change in global activation.

Consistent with the notion that correlation is invariant to mean levels of activation, we found that the *uniform* activation increase yielded no differences in correlations across pair types, regardless of the underlying correlation between stimuli (**Figure 2a**). However, consistent with our previous work highlighting that observed correlations index the proportion of shared (relative to total) across-voxel variability between events (Davis et al., 2014), an increase in activation that was not uniform across voxels *did* produce differences in correlation values across pair types. Specifically, relative to SN pairs, the *subset* activation increase yielded a selective increase in SS pair correlations when the underlying stimulus patterns were uncorrelated, and an additional smaller increase in NN pair correlations when the underlying stimulus patterns were correlated (**Figure 2b**). This is because the additional activation added shared variability to the SS pairs, and, when the underlying correlation was greater than zero, also added detrimental unshared variability to the SN pairs. The *amplified* increase in activation yielded no differences in correlations across conditions when the underlying stimulus patterns were uncorrelated, but yielded an increase in SS pair correlations and a decrease in NN pair correlations (relative to SN pairs) when the underlying stimulus patterns were correlated (**Figure 2c**). Here, the additional activation increased the ability to detect pre-existing shared variability and brought the observed correlation closer to the true correlation. The SS pairs benefitted most from this increase in the ability to detect shared variance (from the two scary stimuli), but the SN pairs also benefitted from the increased ability to detect shared variability (from the one scary stimulus). In all cases, these effects became stronger as the magnitude of global activation increased.

**Figure 2.**
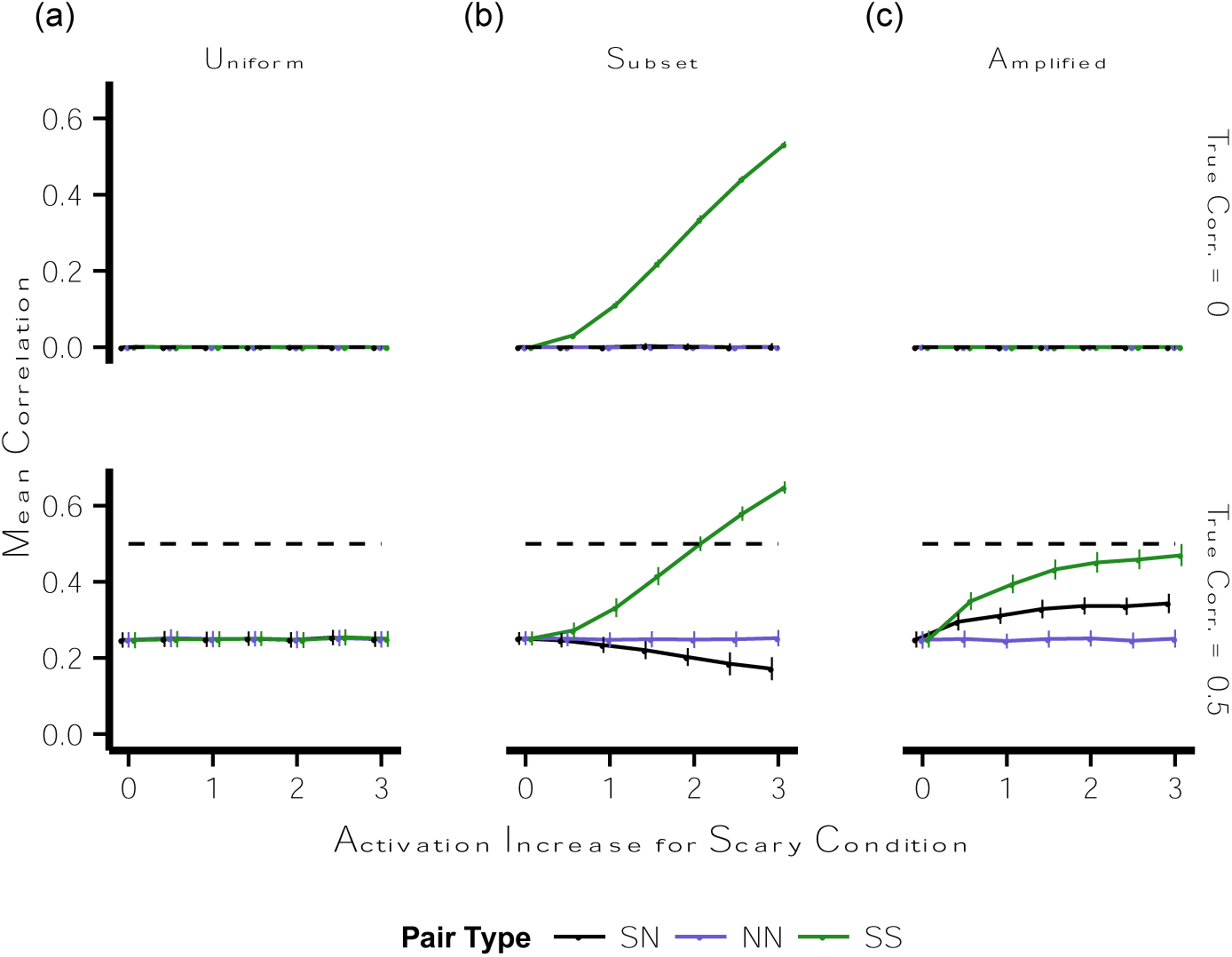
Effects of increases in activation on multivariate pattern similarity. Points indicate the mean observed correlation for pairs of stimuli at parametrically varying levels of an increase in activation for stimuli in the scary condition; note that 0 on the x-axis indicates no increase in activation. Error bars are standard deviation across 100 simulations. Dashed lines indicate ground-truth levels of correlation (which do not vary by condition). *Left-right*: activation was added to stimuli in the scary condition as **(a)** a uniform increase across all voxels (*uniform* case), **(b)** a uniform increase for 50% of the voxels (*subset* case), or **(c)** a stimulus-specific increase across all voxels equal to a multiple of the baseline response to each stimulus (*amplified* case). ***Top-bottom***: underlying correlation across voxels was set to .00 or .50. *NN* = pairs of stimuli both drawn from the non-scary condition, *SS* = pairs of stimuli both drawn from the scary condition, *SN* = pairs of stimuli in which one was drawn from the scary condition and one was drawn from the non-scary condition, *corr.* = correlation.

### Can we statistically correct for the influence of global activation on pattern similarity?

We next asked whether it is possible to statistically correct for the effects observed in the previous simulations. In other words, can we use statistical adjustments to obtain estimates of pattern similarity that are independent of global increases in activation? To do so, we used our simulation framework to examine whether traditional statistical adjustment techniques are able to recover a null pattern similarity effect in the face of our *subset* and *amplified* activation effects. Because the *amplified* simulation relies on a condition-independent correlated pattern of response across voxels, the covariance parameter for all simulations was set to .50 (i.e., ground-truth across-voxel patterns had a mean correlation of .50). Simulations in which the covariance parameter was set to .00 (i.e., ground-truth across-voxel patterns had a mean correlation of .00) can be found in the **Supplementary Materials**.

### Approach one: partial out activation after pattern similarity has been computed

One family of statistical approaches to mitigate the influence of effects of activation on pattern similarity is to calculate correlation and activation metrics using the observed data and then control for activation at the final level of the analysis, i.e., to partial out the effect of activation from one or more of the variables of interest before examining how these variables relate to each other. Critically, the success of this family of approaches hinges on the degree to which the relationship between activation and correlation is successfully captured by the model used to link these measurements.

To examine the efficacy of this approach in recovering ground-truth correlations, we simulated data using the *subset* and *amplified* simulations previously described. For each pair of stimuli we calculated (a) the observed correlation between the stimuli and (b) the sum of the activation (mean activation across voxels for a given stimulus) of the two stimuli. We regressed correlations on activation and then examined whether the residuals from this regression differed by pair type. Because this procedure will remove not only effects related to activation but also the intercept, i.e., the mean correlation across stimulus pairs, we focus on whether the pattern of data yielded by this procedure reflects the *qualitative* rather than *quantitative* pattern of ground-truth correlations.

The results of this statistical adjustment are shown in **Figure 3-top** (see **Figure S1** for results from the *uniform* simulation and simulations in which the covariance parameter was set to .00). In the case of the *subset* simulation this procedure created a new pattern of data that differed both from the ground-truth data and from the observed data: NN and SS correlations were now equivalent, and both were greater than SN correlations. The reason for this becomes apparent when looking at the relationship between correlation and activation from a single representative simulation (**Figure 5a**): the relationship between correlation and activation is not well modeled by the linear function used in the regression and the residuals from this model reflect this poor fit. In the case of the *amplified* simulations this procedure largely eliminated the previously observed condition-specific differences in observed correlations. Again, data from a single representative simulation (**Figure 5a**) reveal that this is due to the relatively linear relationship between correlation and activation. Importantly, even in the *amplified* case, the use of a linear function will no longer be successful if correlation does not scale linearly with activation; we can expect the linearity of this relationship to break down when correlations become bounded by floor or ceiling effects, such as under conditions of high noise or high signal. Finally, although for simplicity we used an increase in activation that was constant across all stimuli within the scary condition, additional simulations confirmed that similar results were obtained when the magnitude of the signal increase was allowed to vary across scary stimuli (inducing a relationship between activation and correlation *within* a given pair type rather than simply a coarse relationship between activation and correlation across pair types; **Figure S2**).

**Figure 3.**
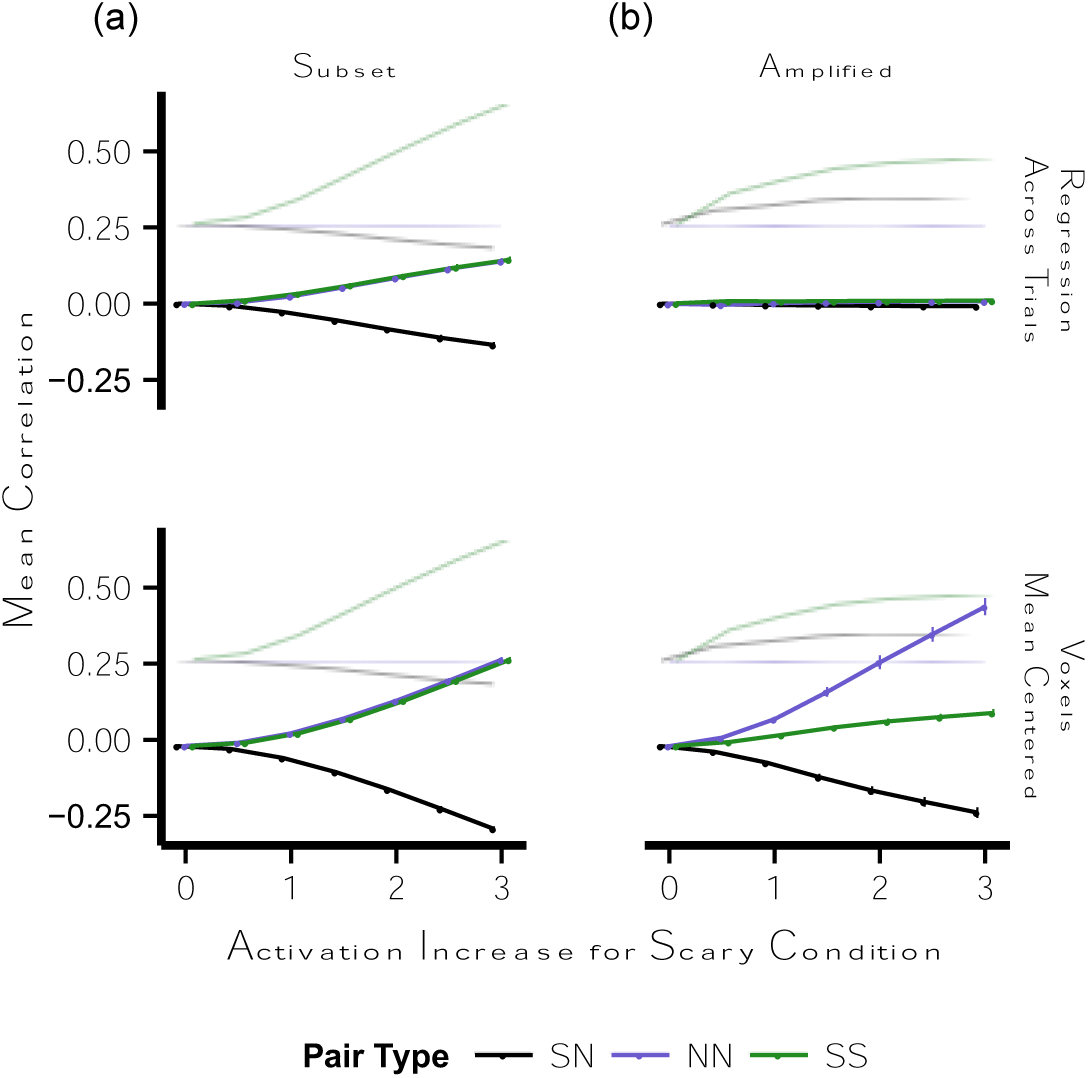
Multivariate pattern similarity following regression adjustments for the effect of activation. Points indicate mean adjusted correlation estimate for pairs of stimuli at parametrically varying levels of an increase in activation for stimuli in the scary condition; note that 0 on the x-axis indicates no increase in activation. Error bars are standard deviation across 100 simulations. Light traces indicate mean observed (unadjusted) correlation. *Left-right*: activation was added to stimuli in the scary condition as either **(a)** a uniform increase for 50% of the voxels (*subset* case), or **(b)** a stimulus-specific increase across all voxels equal to a multiple of the baseline response to each stimulus (*amplified* case). ***Top-bottom***: (top) residuals after regressing correlation on activation across all pairs of stimuli; (bottom) correlations between residuals after mean-centering each voxel. *NN* = pairs of stimuli both drawn from the non-scary condition, *SS* = pairs of stimuli both drawn from the scary condition, *SN* = pairs of stimuli in which one was drawn from the scary condition and one was drawn from the non-scary condition.

These results confirm that attempts to partial out the effect of activation from correlation estimates can successfully recover the true underlying relationship between correlation and condition only if the model used to link activation and correlation successfully captures the relationship between these two measurements. The success of this link depends on the function used to compute a single measure of activation across two distinct stimuli (e.g., additive, multiplicative, maximum) and on the function used to relate correlation and activation (e.g., linear, quadratic), both of which are difficult to specify a priori.

### Approach two: partial out activation before pattern similarity has been computed

A second family of statistical approaches to mitigate the influence of activation on pattern similarity is to attempt to control for mean activation on a voxel-wise basis *before* calculating pattern similarity across voxels. Critically, the success of this family of approaches hinges on the degree to which unwanted effects of activation can be isolated and removed at the level of the individual voxel.

To examine the efficacy of this approach in recovering ground-truth correlations, we again simulated data using the *subset* and *amplified* simulations described above. We then centered each voxel by subtracting the mean activation of that voxel across all stimuli from that voxel’s response to each stimulus, computed correlations across the remaining values in each voxel, and examined whether these correlations differed by pair type.

This statistical adjustment was not successful in recovering the qualitative pattern of ground-truth correlations in either the *subset* or *amplified* case (**Figure 3-bottom;** see **Figure S1** for results from the *uniform* simulation and simulations in which the covariance parameter was set to .00). Specifically, mean centering failed to isolate and remove the critical activation component driving changes in the patterns of activity across voxels. In the *subset* case, the mean response in each voxel incorporates both the magnitude of the response to non-scary stimuli and to scary stimuli, and thus removing the mean for a given voxel only removes *part* of the activation that was added to scary stimuli, and removes *baseline* activation from the non-scary stimuli. In the *amplified* case, the increase in pattern similarity is driven by a unique pattern of activation that is specific to each voxel-by-stimulus pair, and thus attempts to isolate and remove this activation by subtracting a *single* constant across all stimuli were unsuccessful.

In failing to recover the qualitative pattern of ground-truth correlations, these statistical adjustments produced new, undesirable differences in correlations across pair types. The precise pattern of results reflects several co-occurring processes (see **Figure 5b** for a representative single simulation). First, mean centering breaks up pre-existing ground-truth covariance across all stimuli, so correlations move toward zero (even in the case in which no activation was added to the scary stimuli). Second, mean centering further *decreases* across-condition (SN) correlations because consistent responses across voxels that were previously shared across condition (but were of different magnitudes) become anti-correlated with respect to condition. Third, only *part* of the activation, and thus only part of the shared across-voxel variability, added to the scary stimuli is removed, thus attenuating, but not eliminating, inflated SS correlations. Finally, *new* shared across-voxel variability is added to the non-scary stimuli, inflating NN correlations. Additional simulations confirmed that similar results were obtained when the magnitude of the activation increase was allowed to vary across stimuli (**Figure S2**).

To summarize, attempts to partial out activation from individual voxels before pattern similarity was computed were unsuccessful. The effects for each given pair type depended upon whether removing the mean from each voxel impacted shared or unshared across-voxel variability. Critically, in no instance did this approach serve to recover the qualitative pattern of pre-existing ground-truth correlations across conditions, and instead produced entirely new patterns of data.

### Approach three: match activation across conditions

A third family of statistical approaches to mitigate the influence of activation on pattern similarity is to remove voxels or stimuli in order to match activation across conditions. Critically, the success of this family of approaches hinges on the degree to which unwanted effects of activation are confined to a limited number of identifiable individual voxels or identifiable individual stimuli.

To examine the efficacy of this approach in recovering ground-truth correlations, we simulated data using the *subset* and *amplified* simulations but modified the simulations to allow for sufficient variability in activation across scary stimuli to perform matching procedures. We then either removed individual voxels that showed differential activation for scary versus non-scary stimuli (identified at either p < .05 or p < .50) or subsampled stimuli such that each remaining scary stimulus was matched in activation to a specific non-scary stimulus. We then computed mean correlations for each pair type using only the remaining voxels or the remaining stimuli.

The results of the voxel removal procedure are displayed in **Figure 4** (see **Figure S3** for results from the *uniform* simulation and simulations in which the covariance parameter was set to .00). Theoretically, this adjustment should be especially useful in attenuating increases in pattern similarity that arise from the *subset* case: selectively eliminating the subset of voxels that show an activation increase for the scary stimuli should recover the ground-truth correlations. Some success for this approach can indeed be seen in the *subset* case when voxels were removed if they discriminated between conditions at p < .05 (see **Figure 5c** for a representative single simulation). However, there is a small trend in which both SS and NN correlations are less than SN correlations, and this pattern is accentuated when voxels were removed if they discriminated between conditions at p < .50. This new pattern arises when voxels that are not part of the problematic subset are removed. Specifically, individual voxels will vary in activation levels, with voxels with more extreme activation exerting a greater influence on correlation values relative to voxels with more central activation values; the removal of voxels that have *consistent* low or high activation levels for one condition will selectively lower the corresponding within-condition correlations. In the *subset* case, the remaining ‘non-subset’ voxels do not systematically differ as a population in terms of activation for scary and non-scary stimuli. In this case, voxels that exhibit consistently high or low activation for scary *or* non-scary stimuli are removed, thus removing shared across-voxel variability from both SS and NN pairs while removing unshared across-voxel variability from SN pairs. A similar principle applies to the *amplified* case. Here, however, there is no ‘subset’ of voxels that can be selectively removed. Instead, voxels that show consistently high activation across conditions become the voxels that show the greatest difference between conditions (because the difference is a multiple of the baseline response); when these voxels are removed, all correlations are attenuated, with the strongest attenuation in SS correlations (see **Figure 5c** for a representative single simulation).

**Figure 4.**
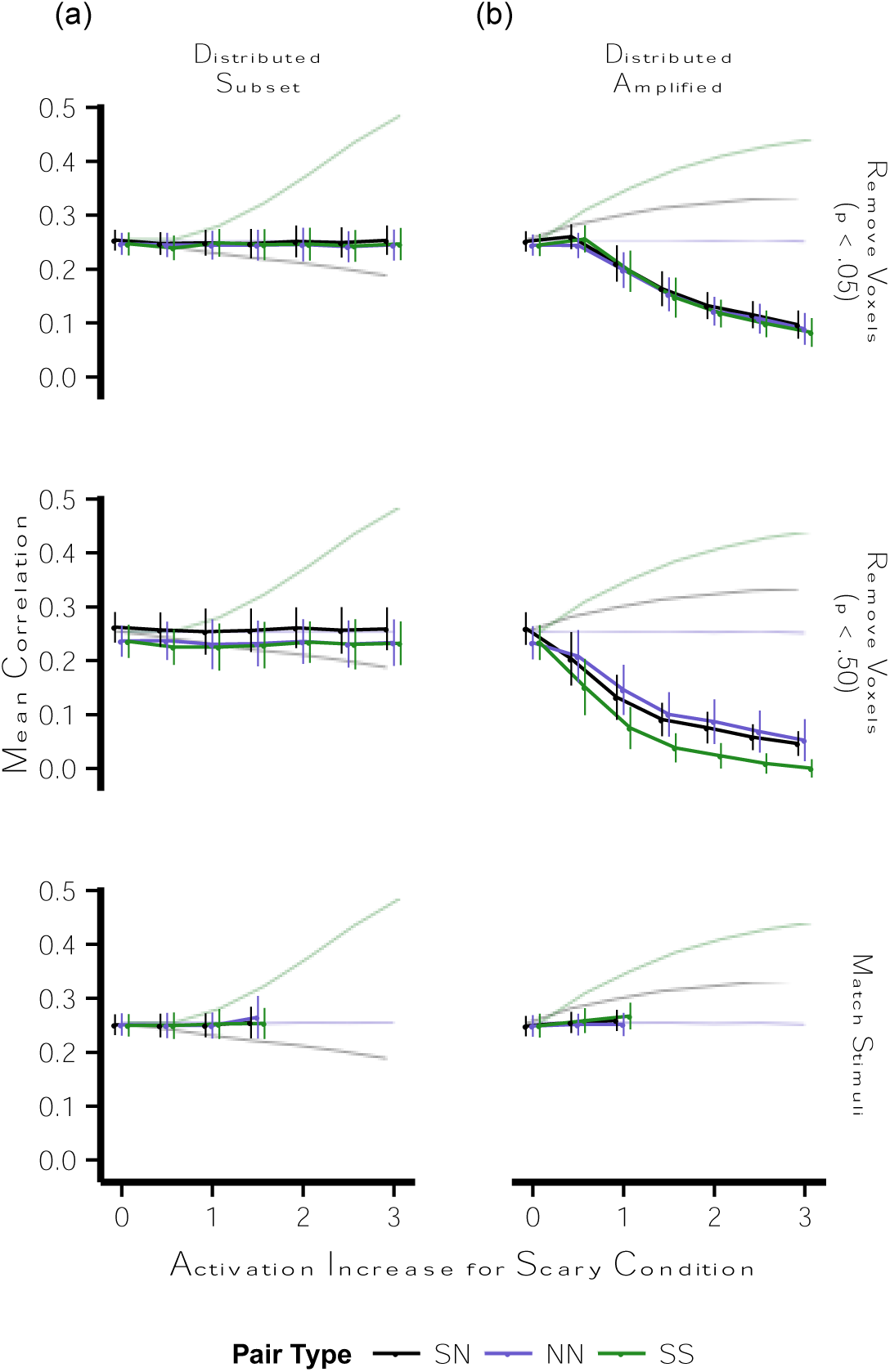
Multivariate pattern similarity following subsampling to match on activation. Points indicate mean adjusted correlation estimate for pairs of stimuli at parametrically varying levels of an increase in activation for stimuli in the scary condition; note that 0 on the x-axis indicates no increase in activation. The magnitude of this increase in activation was distributed across stimuli according to a χ^2^ distribution with a parametrically varying mean. Error bars are standard deviation across 100 simulations. Light traces indicate mean observed (unadjusted) correlation. If fewer than 5 of the 100 simulations (subsampling attempts) were successful for a given combination of parameters, the results for this combination of parameters are not included. *Left-right*: activation was added to stimuli in the scary condition as either **(a)** a uniform increase for 50% of the voxels (*subset* case), or **(b)** a stimulus-specific increase across all voxels equal to a multiple of the baseline response to each stimulus (*amplified* case). ***Top-bottom***: (top) observed correlations after removing voxels showing a significant difference in activation between scary and non-scary stimuli at p < .05; (middle) observed correlations after removing voxels showing a significant difference in activation between scary and non-scary stimuli at p < .50; (bottom) observed correlations after retaining only a subset of stimuli that are matched on mean activation across voxels. *NN* = pairs of stimuli both drawn from the non-scary condition, *SS* = pairs of stimuli both drawn from the scary condition, *SN* = pairs of stimuli in which one was drawn from the scary condition and one was drawn from the non-scary condition.

The stimulus matching procedure attenuated, but did not always completely eliminate, pattern similarity differences across pair types (**Figure 4**); the success of the procedure diminished as the difference in activation between the two conditions grew, presumably because the matching procedure grew less successful (note that in our simulation framework the activation increase in the *subset* condition is half that of the *amplified* condition). Specifically, if the two conditions are roughly matched on activation but still differ systematically such that stimuli from one condition have higher activation than stimuli from the other condition or such that the distribution of activation values differs across condition, the matching procedure will not be successful. In practice, however, the success of the matching procedure can be checked using balance metrics before computing pattern similarity measures (see **Figure 5d** for representative single simulations).

**Figure 5.**
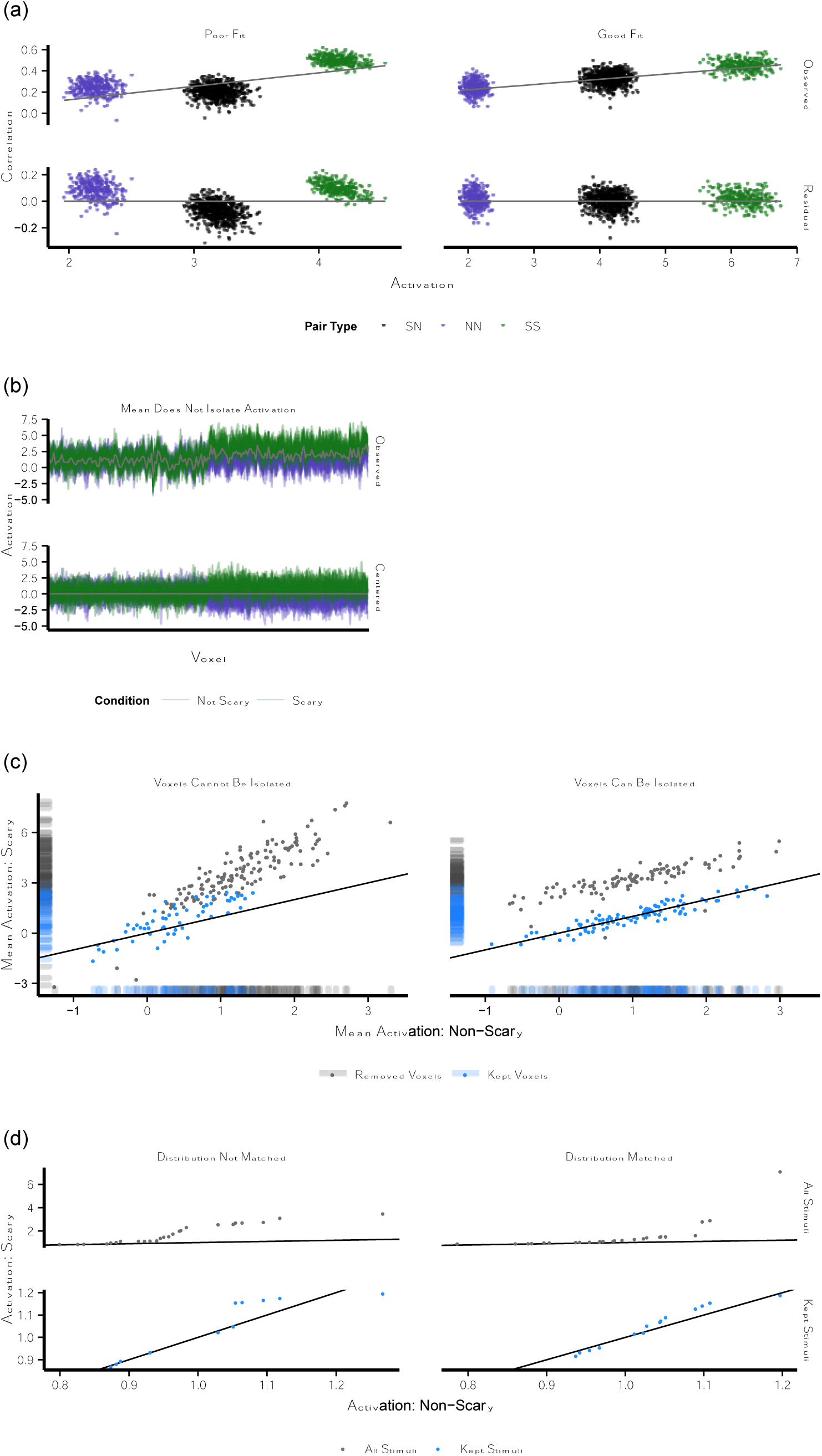
Representative single simulations illustrating failed and successful statistical adjustments for the influence of global activation on pattern similarity. All simulations had an activation increase parameter of 2, with the exception of the stimulus subsampling simulation (panel (d)), which had an activation increase parameter of 1 (matching on activation was not possible with an activation increase parameter of 2). **(a)** Observed correlations (*top*) and residual correlations after regressing observed correlation on activation across all pairs of stimuli (*bottom*) in (left) a case in which the regression function does not fit the data well (here, a representative *subset* case) and (right) a case in which the regression function fits the data well (here, a representative *amplified* case). Points indicate all possible pairs of stimuli; line indicates the best-fit linear regression line relating activation and correlation. When the regression function does not capture the relationship between activation and correlation, a new pattern of results emerges in the residuals; when the regression function does capture the relationship between activation and correlation, the residuals reflect the qualitative pattern of ground-truth correlations (no difference in correlation across pair type). **(b)** Observed (*top*) and mean-centered (*bottom*) across-voxel response to each stimulus in a representative *subset* case. Light traces indicate the response of each voxel to individual stimuli; gray line indicates the mean response of each voxel across all stimuli. Mean centering removes across-voxel variability that is shared across all stimuli, only removes some of the shared across-voxel variability that is selective to the scary stimuli, and adds new shared across-voxel variability to the non-scary stimuli. **(c)** Mean voxel-wise responses to scary and non-scary stimuli after subsampling voxels to match on mean activation (i.e., after removing voxels that respond differentially to the two conditions at p < .05) in (left) a case in which voxels contributing to the increase in SS correlations cannot be isolated (here, a representative *amplified* case) and (right) a case in which voxels contributing to the increase in SS correlations can be isolated (here, a representative *subset* case). Points indicate individual voxels that were either kept or removed; line indicates identity line such that mean activation is equal for scary and non-scary stimuli. Voxels that are far from the identity line and have low variance across stimuli within conditions are most likely to be removed. When voxels inflating SS correlations cannot be isolated, voxels that increase correlations within a given condition (voxels that have relatively high or low activation for that condition and have low variability within that condition) and voxels that contribute to low across-condition correlations (voxels that consistently differ in activation across the two conditions) are removed, decreasing within-condition correlations and increasing across-condition correlations; when voxels inflating SS correlations can be isolated, few voxels outside of this subset of voxels are removed and the pattern of ground-truth correlation can be recovered. **(d)** Quantile-quantile plots of activation for scary and non-scary stimuli before (*top*) and after (*bottom*) subsampling stimuli to match on mean activation in (left) a case in which the distributions of activation for the subsampled scary and non-scary stimuli are not well matched (here, a representative *amplified* case) and (right) a case in which the distributions of activation for the subsampled scary and non-scary stimuli are well matched (here, a representative *subset* case). Points indicate pairs of scary and non-scary stimuli, ordered by activation within each condition; line indicates identity line such that activation is equal for the n^th^ ordered stimulus in the two conditions. Points that fall far from the line indicate differences in the distributions of activation across the two conditions, even if the mean of the distributions (here, level of activation) are well matched. The degree to which the subsampled stimuli in the two conditions share the same distribution will influence how effectively subsampling to match stimuli on activation will recover the pattern of ground-truth correlations. *NN* = pairs of stimuli both drawn from the non-scary condition, *SS* = pairs of stimuli both drawn from the scary condition, *SN* = pairs of stimuli in which one was drawn from the scary condition and one was drawn from the non-scary condition.

To summarize, the success of recovering ground-truth correlations by removing individual voxels that showed an effect of condition on activation depended on how well such a subset of voxels could be isolated and removed. For example, this procedure worked well in the *subset* case using a threshold of p < .05 but not using a threshold of p < .50. Unfortunately, it is challenging to know a priori what the ‘right’ threshold is, as this will vary with the signal properties of a given experiment. In practice, however, visual and/or automated inspection of the data (e.g., histograms of activation differences between conditions across voxels) may ultimately prove useful in determining whether this procedure is appropriate and, if so, guiding an appropriate choice of threshold. The success of attempts to recover ground-truth correlations by retaining a subset of stimuli matched on activation depended on the success of the matching procedure, highlighting the importance of balance metrics in evaluating the success of the matching procedure and in determining whether matching is feasible for a given data set.

### Approach four: include a control condition matched on activation

The final approach that we examined was the inclusion of a planned control condition in the experimental design. Specifically, we looked at the inclusion of additional control cells that are matched with the cells of interest on their levels of a psychological dimension (and thus, theoretically, levels of activation), but in which one does not expect to see the pattern similarity effect of interest. An interaction, such that a pattern similarity effect is selective to, or greater for, the cells of interest could argue against the possibility that differences in activation across levels of this psychological dimension generated the observed pattern similarity effect. However, as we demonstrate below, the success of this approach relies on similar baseline levels of pattern similarity across the control and experimental conditions.

We simulated data mimicking an extension of the design used in the simulations above. Specifically, we simulated a design in which each scary or non-scary stimulus is presented multiple times. Based on a hypothesis that a given region performs a transformation on its inputs that changes where stimuli are situated *relative to one another* in a multidimensional space, one would predict that the observed increase in *across-stimulus* pattern similarity (pattern similarity between distinct stimuli) for scary relative to non-scary stimuli would not be observed when examining *within-stimulus* pattern similarity (pattern similarity between repetitions of the same stimulus). We simulated a 2 x 2 design such that across-stimulus and within-stimulus conditions both included scary and non-scary levels: if the above hypothesis is correct, both conditions should show similar effects of scariness on activation, but only the across-stimulus condition should show an effect of scariness on pattern similarity. Thus, the key question of interest is whether there is an interaction such that the effect of scariness on pattern similarity differs for across-stimulus versus within-stimulus pairs.

First, we simulated data in which ground-truth across-stimulus and within-stimulus correlations were of equivalent magnitude (both covariance parameters = .50). In this case, neither the *subset* nor *amplified* case yielded a pattern of data suggestive of the critical interaction: both the across-stimulus and within-stimulus pairs had equivalently higher SS relative to NN correlations (**Figure 6;** see **Figure S4** for results from the *uniform* simulation and simulations in which the magnitude of the activation increase was allowed to vary across stimuli). This pattern of results would suggest that *differences in activation* between scary and non-scary stimuli (rather than a change in how stimuli are situated relative to one another in a multidimensional space) yielded pattern similarity differences between scary and non-scary stimuli.

**Figure 6.**
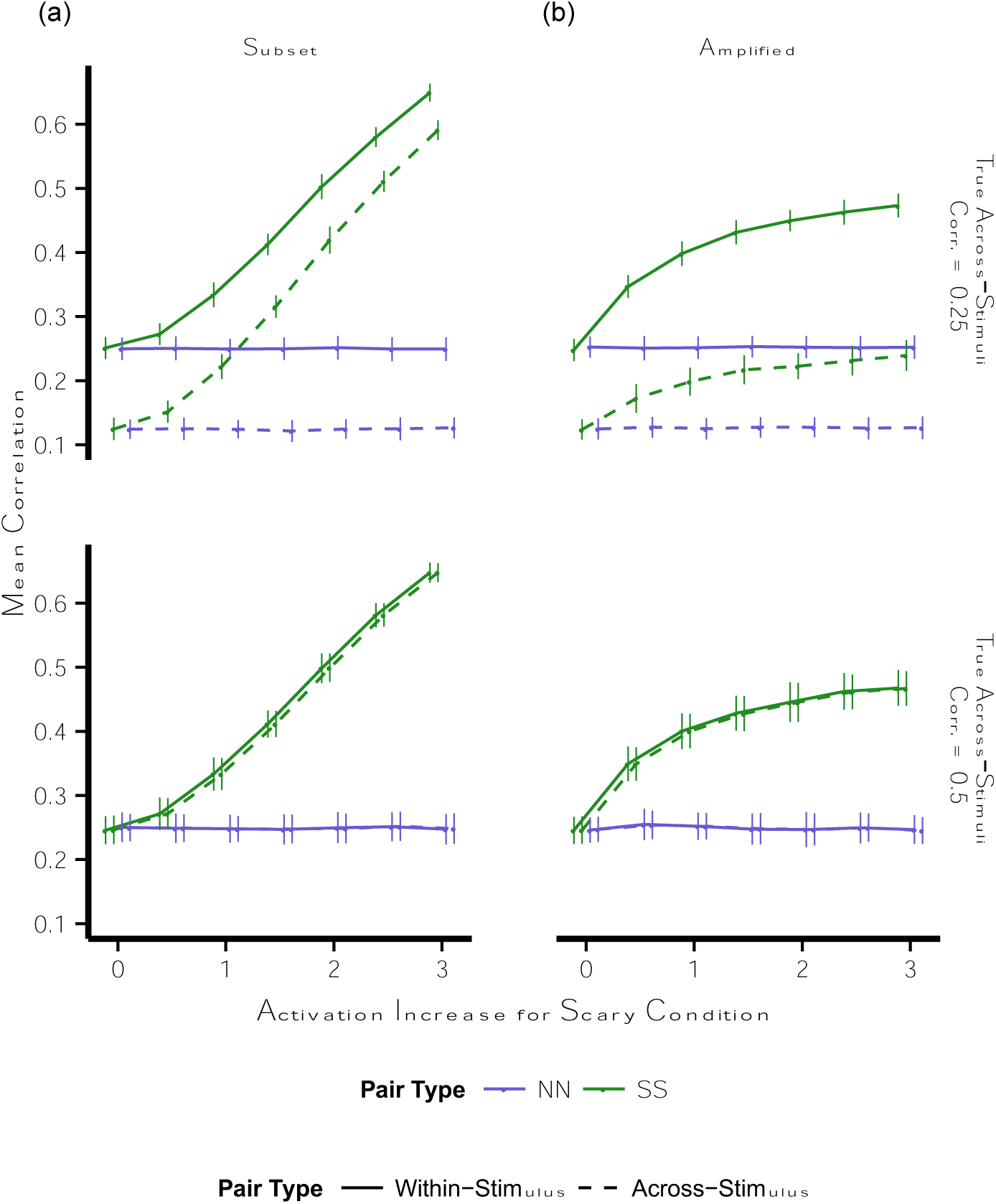
Multivariate pattern similarity in a design including experimental and control conditions. Points indicate mean adjusted correlation estimate for pairs of stimuli at parametrically varying levels of an increase in activation for stimuli in the scary condition; note that 0 on the x-axis indicates no increase in activation. Error bars are standard deviation across 100 simulations. Activation was added to stimuli in the scary condition as either **(a)** a uniform increase for 50% of the voxels (*subset* case), or **(b)** a stimulus-specific increase across all voxels equal to a multiple of the baseline response to each stimulus (*amplified* case). ***Top-bottom***: (top) underlying covariance for distinct stimuli across voxels was set to .25 (ground-truth within-stimulus correlation greater than across-stimulus correlation) or (bottom) .50 (ground-truth within-stimulus correlation same as across-stimulus correlation). *NN* = pairs of stimuli both drawn from the non-scary condition, *SS* = pairs of stimuli both drawn from the scary condition, *corr.* = correlation.

However, this approach was not successful (i.e., the data yielded the critical interaction) when we simulated data in which ground-truth within-stimulus correlations were greater than ground-truth across-stimulus correlations (covariance parameters for within-stimulus and across-stimulus pairs were .50 and .25, respectively; **Figure 6;** see **Figure S4** for results from the *uniform* simulation and simulations in which the magnitude of the activation increase was allowed to vary across stimuli). Specifically, in the *subset* case, the magnitude of the increase in pattern similarity for SS relative to NN pairs was greater for across-stimulus pairs relative to within-stimulus pairs. In this case, the increase in activation adds new shared across-voxel variability to the existing patterns, and the effect of a fixed increase in activation on pattern similarity is nonlinear and depends on pre-existing levels of shared variability. Here, the same increase in activation led to a greater increase in pattern similarity for across-stimulus SS pairs than within-stimulus SS pairs because pre-existing shared variability was lower for across-stimulus relative to within-stimulus SS pairs. Conversely, in the *amplified* case, the magnitude of the increase in pattern similarity for SS relative to NN pairs was smaller for across-stimulus pairs relative to within-stimulus pairs. Although this interaction was in the opposite direction as hypothesized in the current framework, it could be interpreted as evidence for a different mechanism of interest, such as selective sharpening of individual stimulus representations for scary stimuli. In this case, the increase in activation allows for the ability to detect pre-existing shared across-voxel variability, and, again, the effect of a fixed increase in activation on pattern similarity is nonlinear and depends on pre-existing ability to detect shared variability. Here, the same increase in activation led to a greater increase in pattern similarity for within-stimulus SS pairs than across-stimulus SS pairs because pre-existing observed pattern similarity was farther from the ground-truth pattern similarity for within-stimulus relative to across-stimulus pairs.

To summarize, attempts to include a planned control condition in the experimental design to mitigate concerns about influences of activation on pattern similarity relies on the assumption that ground-truth correlations are matched between the experimental and control conditions. When this is not the case, interactions may emerge that do not reflect the hypothesized selectivity of a pattern similarity effect to a given condition.

## Discussion

The present work demonstrates two ways in which differences in global activation across conditions can produce condition-specific differences in pattern similarity measures. Moreover, it suggests that the ability to statistically adjust for these differences in global activation is heavily dependent upon signal parameters that will likely vary widely across experiments and in practice are difficult to isolate outside of the simulation framework employed here.

Consistent with the notion that correlations are invariant to mean differences in activation across conditions, we found that pattern similarity was unaffected by activation increases that were uniform across all voxels. However, a condition-specific increase in activation that was restricted to a subpopulation of voxels did yield pattern similarity differences across conditions. Specifically, pattern similarity for stimuli within the condition with higher activation (SS pairs) increased due to added shared across-voxel variability, whereas pattern similarity between stimuli in different conditions (SN pairs) decreased due to added unshared across-voxel variability. Here we focused on a simple scenario in which a region has two distinct subpopulations of condition-responsive and condition-nonresponsive voxels; however, these pattern similarity effects can arise in any situation in which the amount of activation for a given condition is heterogeneous across voxels (Davis et al., 2014).

In addition, we found that pattern similarity was affected by a condition-specific increase in activation that amplified the response in each voxel according to its baseline response to each individual stimulus. While there has been attention to the idea that correlation (and MVPA in general) is sensitive to the degree of signal (here, activation) relative to noise (Smith et al., 2011; Tong et al., 2012), there has been little focus on this sensitivity as it pertains to differences in signal-to-noise ratio across conditions within a single anatomical region (but see LaRocque et al., 2013). Here, we demonstrate that this type of activation increase boosts the ability to detect pre-existing shared across-voxel variability for stimuli in the condition with higher activation (here, scary stimuli), resulting in increased pattern similarity for SS pairs and, to a lesser extent, SN pairs. For simplicity we simulated an increase in signal with no concurrent increase in noise, however, this effect only requires that the increase in signal outweighs any increase in noise.

Although here we used discrete conditions for illustrative purposes, these results extend to any circumstance in which activation varies systematically with experimental effects of interest. For example, a continuous case in which activation varies parametrically along a dimension of interest could produce a similarly graded pattern similarity effect. More broadly, the demonstration that activation impacts pattern similarity has implications for any comparison of pattern similarity across groups of stimuli that differ in activation, such as comparisons of stimuli encountered at different time points within a scan session.

As most of our simulations suggest that increases in activation for a given condition will increase pattern similarity for pairs of stimuli within that condition, one question of interest is whether increases in activation can *decrease* pattern similarity. Although not simulated here, there are several instances in which this could be the case. For example, if an increase in activation manifested as a randomly distributed effect across voxels and stimuli, this effect would add noise to the data and decrease any pre-existing nonzero correlation. Additionally, if the underlying correlation across stimuli were negative, an *amplified* activation increase would move the observed correlations closer to the true negative correlation. A related question is how a *decrease* in activation relative to baseline will affect pattern similarity. In the *subset* case, this would still amount to the addition of meaningful variance, thereby increasing pattern similarity. Conversely, in the *amplified* case a reduction in the ability to detect shared variance would move correlations closer to zero.

The observed relationship between activation and pattern similarity could theoretically be overcome if it were possible to statistically adjust for the influence of activation on pattern similarity and recover an estimate of pattern similarity that is independent of activation. To this aim, we examined the efficacy of several statistical approaches for removing the influence of activation on pattern similarity.

The success of attempts to remove the influence of activation after correlations were already computed was dependent on the degree to which the relationship between activation and correlation was linear. When our linear function was a poor fit to the data, the resulting pattern similarity estimates were qualitatively altered both from the originally observed pattern of data and from the ground-truth pattern of data. Although it is possible that a different function (such as a quadratic function) or a different method of combining activation across the two stimuli in a given pair (such as taking the minimum activation across the two stimuli) may have been more successful in attenuating the relationship between correlation and activation in our simulations, it is virtually impossible to infer these functions a priori. One may also ask if our failure to successfully remove the influence of activation arose from our specific approach, which was akin to a semipartial correlation between correlation and pair type. It is likely that different approaches will be most successful in different situations, and our example here is meant to illustrate the general circumstances under which this family of approaches can succeed or fail. Finally, although not simulated, similar statistical corrections could be done at the participant-level, such that condition-specific differences in correlation and activation for each participant are entered into a regression. The success of this approach will again rely on the extent to which the regression properly models the relationship between activation and correlation. The effect of a fixed change in activation on correlations will depend on a number of signal parameters (such as baseline correlation and noise) that will differ across participants, which may once again prove difficult to model with a single function that is specified a priori.

Attempts to partial out activation at the voxel level by mean centering did not isolate and remove the critical activation components from each voxel and recover ground-truth correlations, but instead qualitatively changed the pattern of the observed data (see also Garrido et al., 2013). First, mean centering broke up pre-existing covariance across stimuli, lowering all correlations. On top of this, the precise effect on each pair type depended on how subtracting a constant (the mean) impacted the degree of shared variability across those pairs of stimuli, but in no case did mean centering recover the true underlying pattern of data.

Attempts to match stimuli from each condition on activation yielded mixed success. Selectively removing voxels that differentiated between conditions yielded initial success at a stringent statistical threshold of p < .05. Importantly, the success for the *subset* case came by selectively removing voxels that showed an effect of condition on activation, whereas success for the *amplified* case was merely a byproduct of the fact that this procedure removed voxels that lowered SS correlations more than they lowered NN correlations. However, when voxels were removed using a more liberal threshold of p < .50, this qualitatively changed the pattern of data by removing voxels that were differentially important for within-condition correlations (a new mechanism in the *subset* case, and an overextension of the existing mechanism in the *amplified* case). Critically, the more stringent threshold was the ‘correct’ threshold here due to specific signal properties of the simulated data set, but in practice the ‘correct’ threshold will vary with each experiment. Given the difficulty in selecting the ‘correct’ statistical threshold for voxel removal, a potential avenue for future work may be to use unsupervised learning approaches to identify and remove subpopulations of voxels that show condition-specific changes in activation. Subsampling stimuli to match activation across conditions was successful when the distribution of activation could be well matched across conditions, but did not fully attenuate the increase in SS correlations when the matching procedure became less successful. Importantly, although here we included all simulations that matched enough stimuli (regardless of the success of the nearest neighbor matching procedure), in practice it is possible to use balance metrics that are independent of pattern similarity measures to assess the success of multiple, more sophisticated matching, thus providing a priori guidance not available using other adjustment approaches (Ho et al., 2011; 2007).

It is important to note that all of the approaches described above rely on measurable differences in activation across conditions. Without such differences, any attempts to correct for, or match on, activation will be fruitless. Situations in which (a) distinct subpopulations of voxels respond with an activation increase to distinct conditions or (b) both positive and negative deflections from baseline activation are scaled will produce similar pattern similarity results but fail to produce any differences in activation across conditions, making them inherently uncorrectable by any of these approaches.

Finally, we attempted to control for activation using a two-factor experimental design. The logic is that a set of control conditions can be matched on activation to the experimental conditions but not yield the same pattern similarity effect, making an interaction a critical indicator of a pattern similarity effect that extends beyond any effect driven by activation alone. We found that this logic held true when the control and experimental conditions were matched on baseline pattern similarity. However, when the control and experimental conditions were not matched on baseline pattern similarity, this control proved ineffective. Specifically, the impact of both the *subset* and *amplified* increases in activation depended on pre-existing levels of pattern similarity, thereby impacting the control and experimental conditions differently and yielding the critical interaction. However, the interactions yielded in these cases were quantitative interactions; the presence of a cross-over interaction may provide better assurance the results of interest are not driven by activation differences across conditions.

Overall, the success of any given statistical adjustment was highly dependent upon signal parameters that cannot be easily observed by the experimenter, such as how effects of activation manifest across voxels and across stimuli. Further complicating this problem is that here we considered two simplified scenarios. In the *subset* case, only some voxels responded to a given manipulation and the nature of the response was uniform across voxels and across stimuli. In the *amplified* case, all voxels responded to a given manipulation and the nature of the response was voxel-by-stimulus specific such that it increased the gain on a pre-existing response. Although here we explored these scenarios separately, they are undoubtedly intermixed in practice. Within a given region, one may observe a combination of responsive and unresponsive voxels, with the responsive voxels showing a mix of uniform, voxel-specific (Davis et al., 2014), and voxel-by-stimulus-specific responses. Thus, there is no easy way to know what ‘case’ one is observing and no general rule about how activation and pattern similarity will relate to one another, further complicating the possibility of selecting the ‘correct’ statistical adjustment for a given set of data. Moreover, although here we simulated data in which there is an effect of condition on activation but no direct effect of condition on pattern similarity, in practice it is likely that the effects of global activation on pattern similarity will be intermixed with direct effects of condition on pattern similarity. Additional simulations (**Supplemental Text** and **Figures S5 and S6**) revealed that condition-specific global increases in activation can amplify, attenuate, eliminate, or reverse direct effects of condition on pattern similarity. Moreover, the statistical adjustments considered here generally do not recover these ground-truth direct effects of condition on pattern similarity when these effects are (a) contaminated by effects of global activation on pattern similarity, (b) paired with an effect of condition on global activation that does not influence pattern similarity (e.g., the *uniform* case), and in some cases (c) when there is no effect of condition on global activation. The exception to this finding was subsampling to match stimuli in each condition on activation, likely because this approach does not rely on any assumptions about the nature of the relationship between activation and correlation.

Even when considering the above challenges, the simplified scenarios examined here do have signatures that may be, in some cases, distinguished from pattern similarity effects arising from normally distributed condition-specific differences across voxels. For example, *subset* responses yield multimodal distributions of activation across voxels, and *amplified* responses yield differences in across-voxel variability across conditions. Improved methods for visualizing the high dimensional space of across-voxel responses within a region may help to identify these signatures when possible. Similarly, comparing the properties of the observed response to those of simulated data generated using the hypothesized response mechanism may also help to identify important deviations between observed and simulated data.

The present simulations explored how activation can affect pattern similarity and the degree to which statistical adjustments can correct for these influences and / or yield unintended consequences. We demonstrate that the relationship between activation and pattern similarity, along with the efficacy of statistical corrections for activation, will vary according to experimental design and participant-specific signal parameters. This variability in the relationship between activation and pattern similarity suggests a need for explicit justification of why a specific adjustment for activation is (or is not) being used during pattern similarity analyses and consideration of how signal parameters will affect the possible outcomes of this adjustment. It also highlights the relative power of approaches, such as subsampling stimuli matched on activation, that make no a priori assumptions about the relationship between activation and pattern similarity. As the use of pattern similarity analyses continues to grow across domains of cognition, from perception to memory to action, the continued development of such statistical techniques for disentangling activation and pattern similarity will be increasingly important for drawing accurate inferences.

## Supplementary Materials

### Supplementary Text

In the main text we consider a case in which there is a change in global activation for scary relative to non-scary stimuli but no additional, independent change in pattern similarity across pair types. An important question is how changes in activation across conditions, and statistical adjustments for these changes, interact with simultaneous, independent changes in pattern similarity across conditions. To do so, we repeated the simulations reported in the main text for two additional sets of parameters: (a) a case in which ground-truth pattern similarity is higher for SS pairs than SN and NN pairs (SS covariance = .50, SN and NN covariance = .25), and (b) a case in which ground-truth pattern similarity is higher for NN pairs than SN and SS pairs (NN covariance = .50, SN and SS covariance = .25). In both cases the effect of activation was such that activation was greater for scary relative to non-scary pairs.

First, we examined the observed pattern of data for the *uniform*, *subset*, and *amplified* cases (with both constant and distributed amounts of activation across stimuli; **Figure S5a,d**). As in the main text, the increase in activation for the scary stimuli did not alter the observed correlations in the *uniform* case. When pattern similarity was higher for SS relative to SN and NN pairs, the increase activation for scary stimuli in the *subset* and *amplified* case magnified this pattern similarity effect; when pattern similarity was higher for NN relative to SN and SS pairs, the increase activation for scary stimuli in the *subset* and *amplified* case attenuated, eliminated, or reversed this pattern similarity effect, depending on the amount of activation in conjunction with the other signal parameters. Note that in the *amplified* case, whether the ground-truth pattern similarity effect of NN > SS can be eliminated or reversed will depend on whether ground-truth SS correlations are of equal or greater magnitude than observed NN correlations.

Next, we assessed the effects of the statistical adjustments reported in the main text on these observed patterns of data (**Figure S5b,c,e,f**). Note that in all cases an activation parameter of zero illustrates an effect of condition on pattern similarity but no effect of condition on activation, and the *uniform* case with an activation parameter greater than zero illustrates independent effects of condition on activation and pattern similarity. In both of these cases, a statistical adjustment that removes changes in correlations that are the byproduct of changes in activation should not change the observed correlations, which are already independent from effects of condition on activation. In all other cases we would want to recover the pattern of correlations that was observed when the activation parameter is set to zero (i.e., when there is no change in activation across condition). The only statistical adjustment that met these criteria was subsampling to match stimuli on activation. Regressing correlation on activation across trials did not change the pattern of data when there was no effect of activation (as the residuals are essentially the observed data with the intercept removed), but did change the qualitative pattern of data in the *uniform* case (as activation and correlation, although independent effects, are related to each other via condition and thus this relationship is quantified and removed from the observed pattern of data). Mean centering and subsampling voxels produced new patterns of data when there was no effect of activation, in the *uniform* case, and in all other cases, suggesting that they are ineffective adjustments for separating activation and correlation.

Finally, we assessed a scenario in which an effect of condition on pattern similarity is present in one set of experimental cells (here, across-stimulus correlations), but not in another set of control cells (here, within-stimulus correlations), and an effect of condition on activation is present in both the experimental and control cells. To do so, we repeated the simulations comparing within-stimulus and across-stimulus pattern similarity reported in the main text for two additional sets of parameters: (a) a case in which ground-truth pattern similarity is higher for across-stimulus SS pairs than across-stimulus NN pairs (across-stimulus SS covariance = .50, across-stimulus NN covariance = .25), and (b) a case in which ground-truth pattern similarity is higher for across-stimulus NN pairs than across-stimulus SS pairs (across-stimulus NN covariance = .50, across-stimulus SS covariance = .25). In both cases the effect of activation was such that activation was greater for scary relative to non-scary pairs, the within-stimulus covariance was .75 for both SS and NN within-stimulus pairs, and the SN covariance was .25. In both of these scenarios one should observe an interaction such that across-stimulus correlations vary between SS and NN across-stimulus pairs but not between SS and NN within-stimulus pairs. Results are displayed in **Figure S6**. The increase in activation for the scary stimuli did not alter the observed correlations in the *uniform* case. However, in the *subset* and *amplified* cases, the increase in activation enhanced, attenuated, eliminated, or reversed the ground-truth interaction in pattern similarity across pair type, depending on the amount of activation in conjunction with the other signal parameters. Again, note that in the *amplified* case, the effect of the increase in activation for the scary condition will depend on the relationship between the ground-truth SS correlations and the observed NN correlations.

### Supplementary Figures

**Figure S1.**
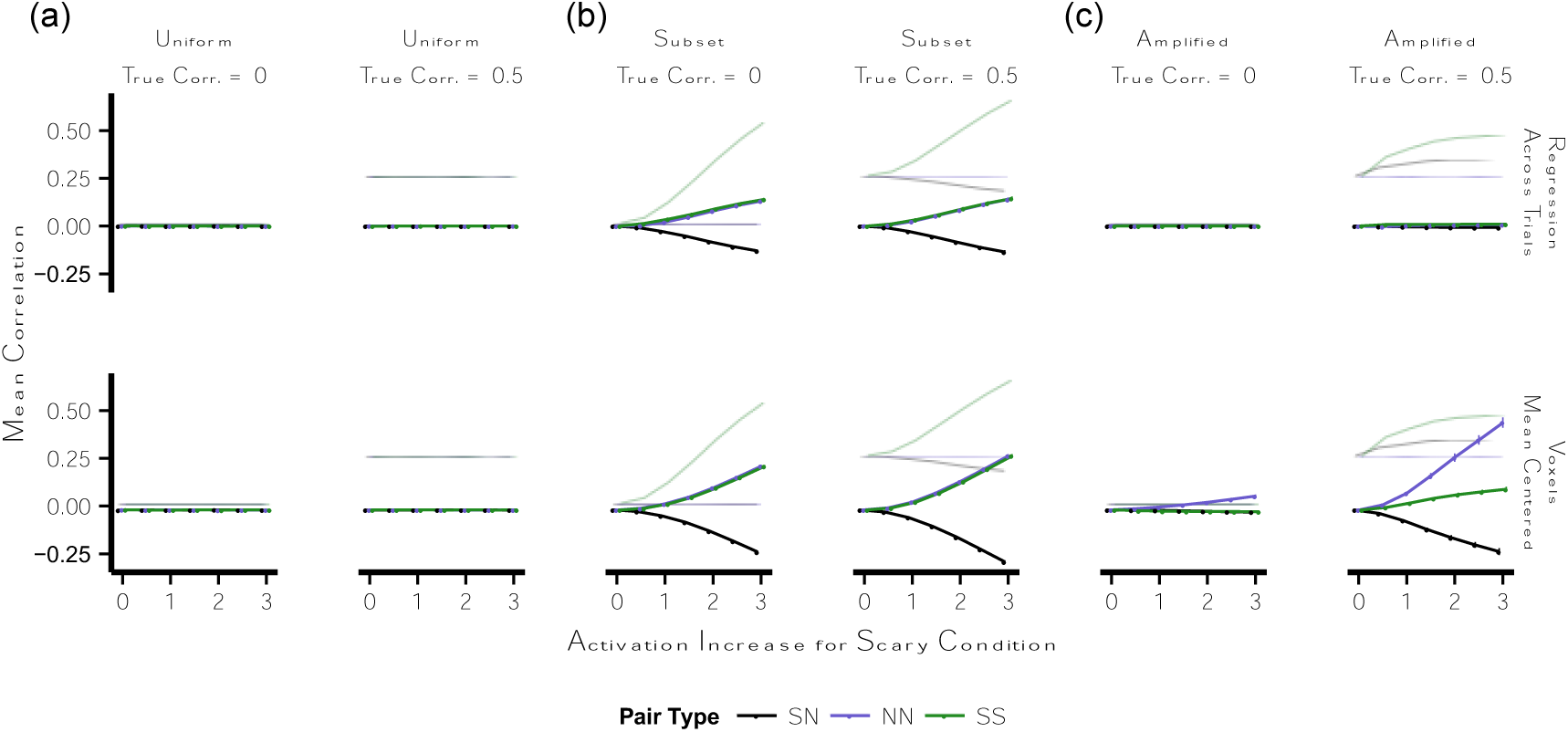
Multivariate pattern similarity following regression adjustments for the effect of activation. Points indicate mean adjusted correlation estimate for pairs of stimuli at parametrically varying levels of an increase in activation for stimuli in the scary condition; note that 0 on the x-axis indicates no increase in activation. Error bars are standard deviation across 100 simulations. Light traces indicate mean observed (unadjusted) correlation. Activation was added to stimuli in the scary condition as **(a)** a uniform increase across all voxels (*uniform* case), **(b)** a uniform increase for 50% of the voxels (*subset* case), or **(c)** a stimulus-specific increase across all voxels equal to a multiple of the baseline response to each stimulus (*amplified* case). Within each type of activation increase, the underlying correlation across stimuli was set to 0.00 or 0.50. ***Top-bottom***: (top) residuals after regressing correlation on activation across all pairs of stimuli; (bottom) correlations between residuals after mean-centering each voxel. *NN* = pairs of stimuli both drawn from the non-scary condition, *SS* = pairs of stimuli both drawn from the scary condition, *SN* = pairs of stimuli in which one was drawn from the scary condition and one was drawn from the non-scary condition, *corr.* = correlation.

**Figure S2.**
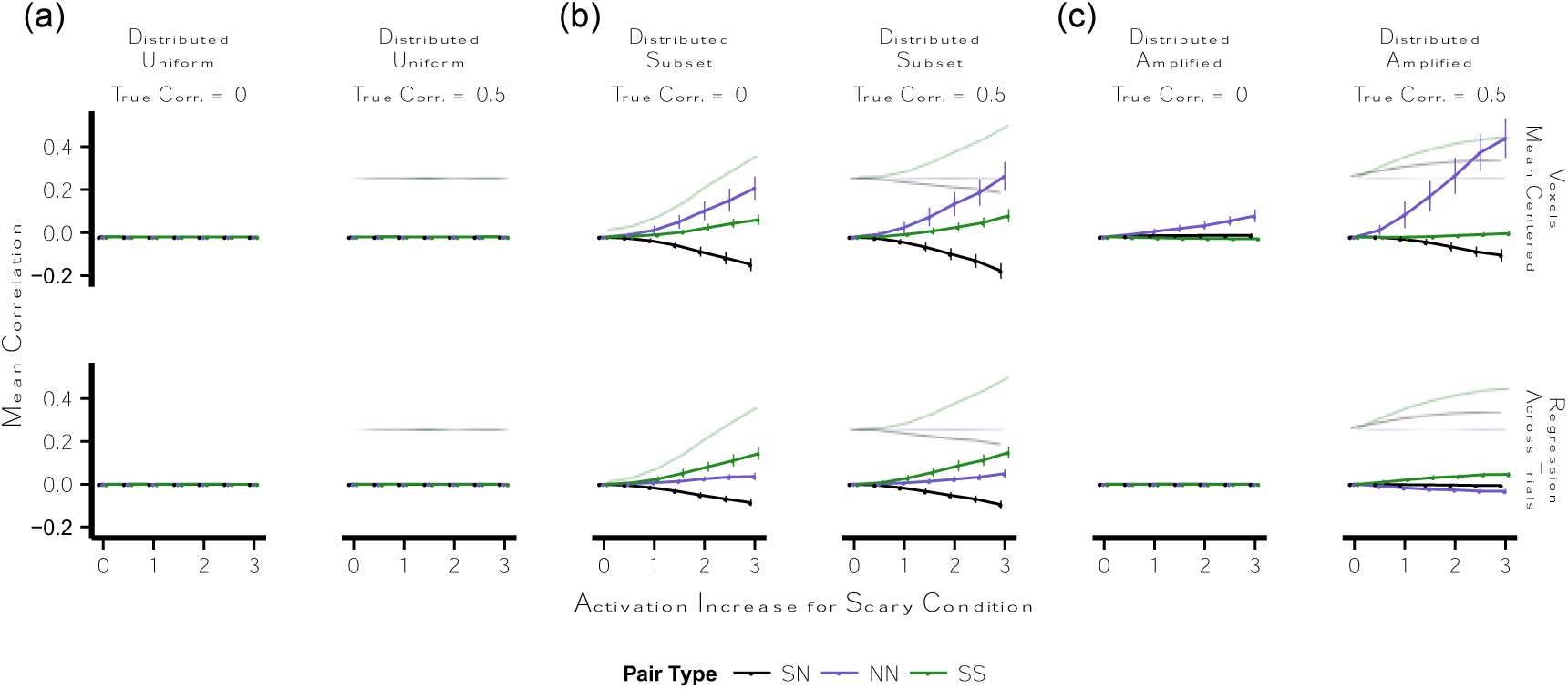
Multivariate pattern similarity following regression adjustments for an effect of activation that is distributed across stimuli. Points indicate mean adjusted correlation estimate for pairs of stimuli at parametrically varying levels of an increase in activation for stimuli in the scary condition; note that 0 on the x-axis indicates no increase in activation. Unlike in the main text, the magnitude of this increase in activation was distributed across stimuli according to a χ^2^ distribution with a parametrically varying mean. Error bars are standard deviation across 100 simulations. Light traces indicate mean observed (unadjusted) correlation. *Left-right*: activation was added to stimuli in the scary condition as **(a)** a uniform increase across all voxels (*uniform* case), **(b)** a uniform increase for 50% of the voxels (*subset* case), or **(c)** a stimulus-specific increase across all voxels equal to a multiple of the baseline response to each stimulus (*amplified* case). Within each type of activation increase, the underlying correlation across stimuli was set to 0.00 or 0.50. ***Top-bottom***: (top) residuals after regressing correlation on activation across all pairs of stimuli; (bottom) correlations between residuals after mean-centering each voxel. *NN* = pairs of stimuli both drawn from the non-scary condition, *SS* = pairs of stimuli both drawn from the scary condition, *SN* = pairs of stimuli in which one was drawn from the scary condition and one was drawn from the non-scary condition, *corr.* = correlation.

**Figure S3.**
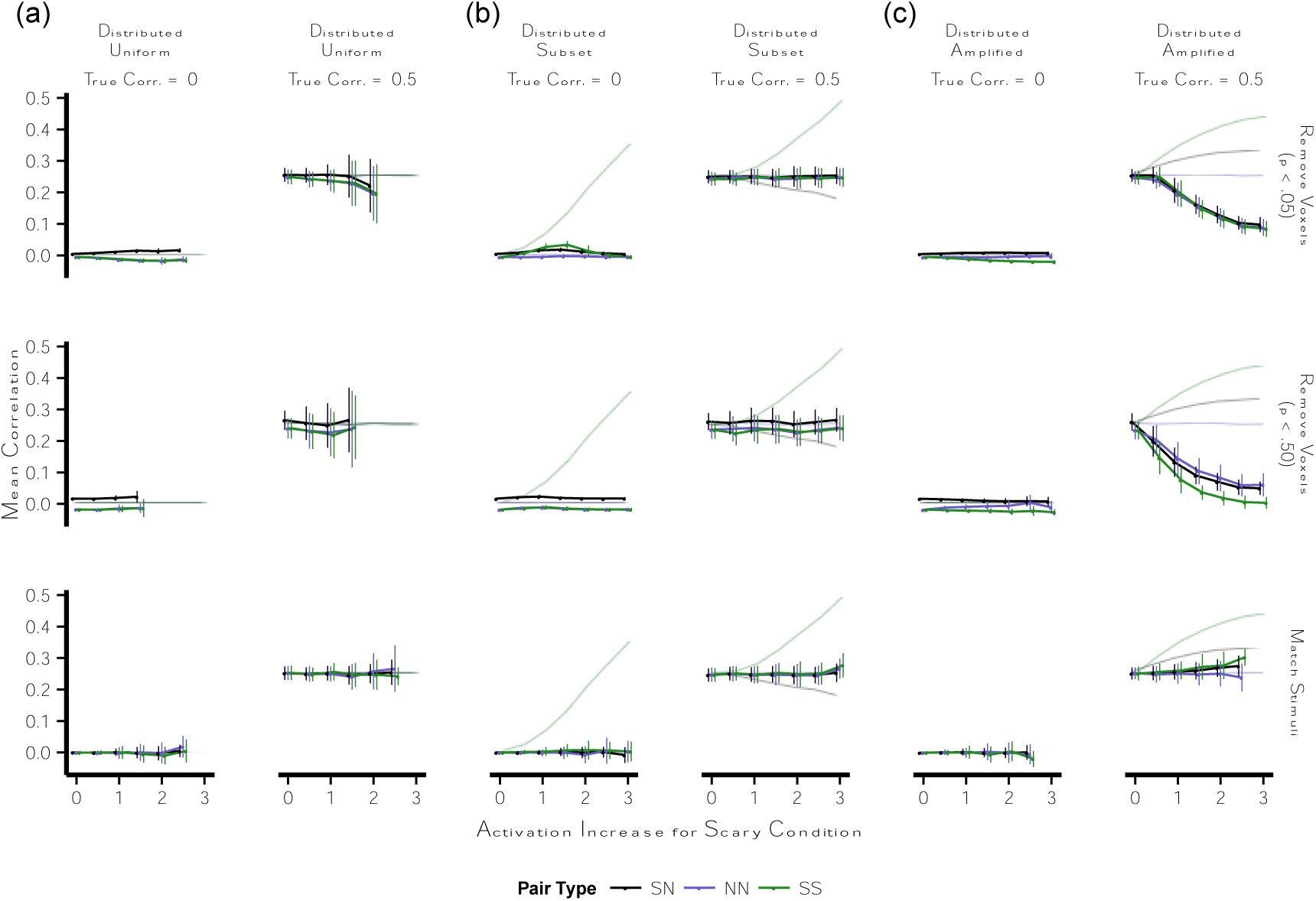
Multivariate pattern similarity following subsampling to match on activation. Points indicate mean adjusted correlation estimate for pairs of stimuli at parametrically varying levels of an increase in activation for stimuli in the scary condition; note that 0 on the x-axis indicates no increase in activation. Error bars are standard deviation across 100 simulations. Light traces indicate mean observed (unadjusted) correlation. If fewer than 5 of the 100 simulations (subsampling attempts) were successful for a given combination of parameters, the results for this combination of parameters are not included. Activation was added to stimuli in the scary condition as **(a)** a uniform increase across all voxels (*uniform* case), **(b)** a uniform increase for 50% of the voxels (*subset* case), or **(c)** a stimulus-specific increase across all voxels equal to a multiple of the baseline response to each stimulus (*amplified* case). Within each type of activation increase, the underlying correlation across stimuli was set to 0.00 or 0.50. ***Top-bottom***: (top) observed correlations after removing voxels showing a significant difference in activation between scary and non-scary stimuli at p < .05; (middle) observed correlations after removing voxels showing a significant difference in activation between scary and non-scary stimuli at p < .50; (bottom) observed correlations after retaining only a subset of stimuli that are matched on mean activation across voxels. *NN* = pairs of stimuli both drawn from the non-scary condition, *SS* = pairs of stimuli both drawn from the scary condition, *SN* = pairs of stimuli in which one was drawn from the scary condition and one was drawn from the non-scary condition, *corr.* = correlation.

**Figure S4.**
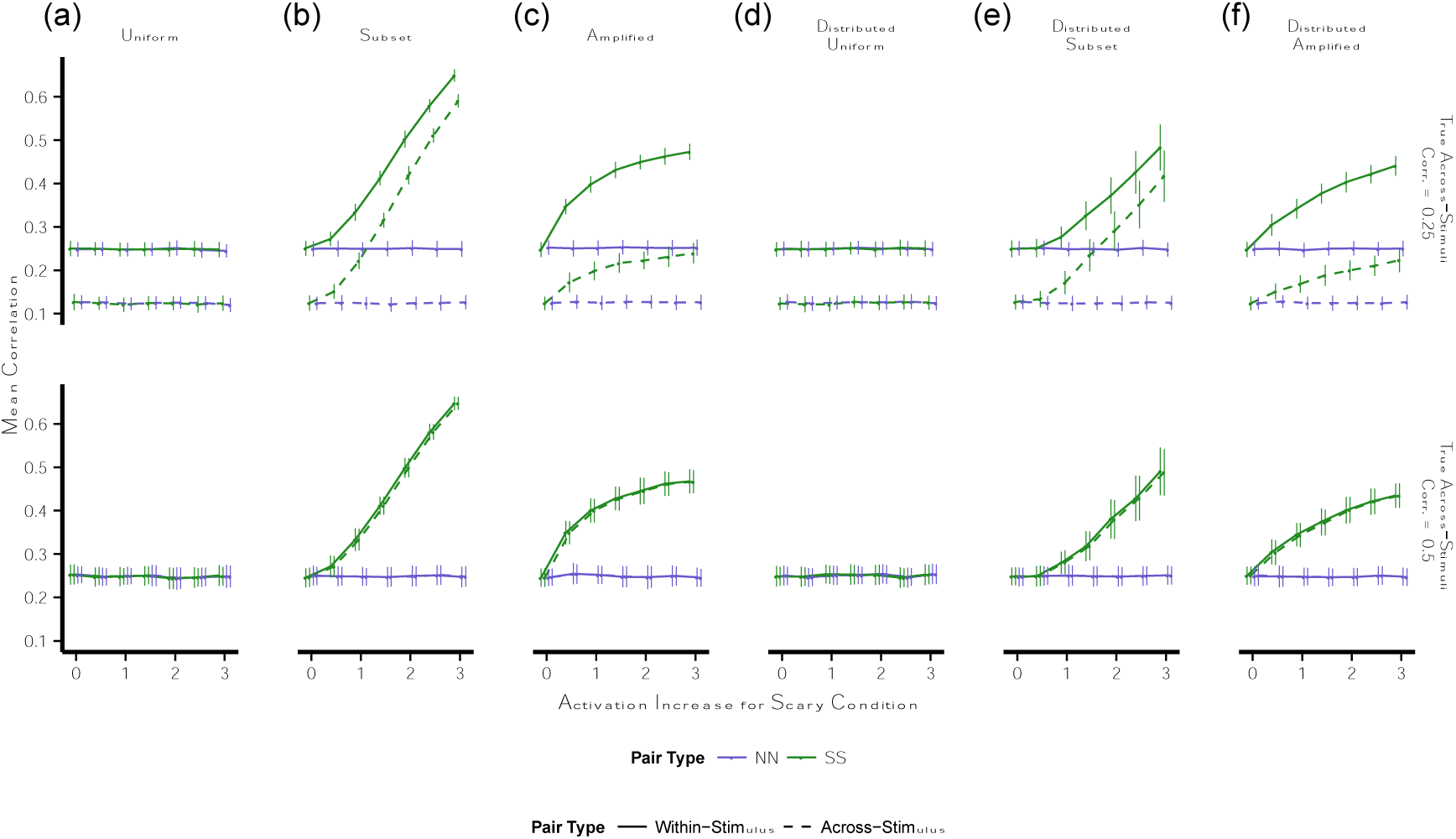
Multivariate pattern similarity in a design including experimental and control conditions. Points indicate mean adjusted correlation estimate for pairs of stimuli at parametrically varying levels of an increase in activation for stimuli in the scary condition; note that 0 on the x-axis indicates no increase in activation. Error bars are standard deviation across 100 simulations. Activation was added to stimuli in the scary condition as **(a, d)** a uniform increase across all voxels (*uniform* case), **(b, e)** a uniform increase for 50% of the voxels (*subset* case), or **(c, f)** a stimulus-specific increase across all voxels equal to a multiple of the baseline response to each stimulus (*amplified* case). Within each type of activation increase, the increase was either (a, b, c) constant across stimuli or (d, e, f) distributed across stimuli according to a χ^2^ distribution with a parametrically varying mean. ***Top-bottom***: (top) underlying covariance for distinct stimuli across voxels was set to .25 (ground-truth within-stimulus correlation greater than across-stimulus correlation) or (bottom) .50 (ground-truth within-stimulus correlation same as across-stimulus correlation). *NN* = pairs of stimuli both drawn from the non-scary condition, *SS* = pairs of stimuli both drawn from the scary condition, *SN* = pairs of stimuli in which one was drawn from the scary condition and one was drawn from the non-scary condition, *corr.* = correlation.

**Figure S5.**
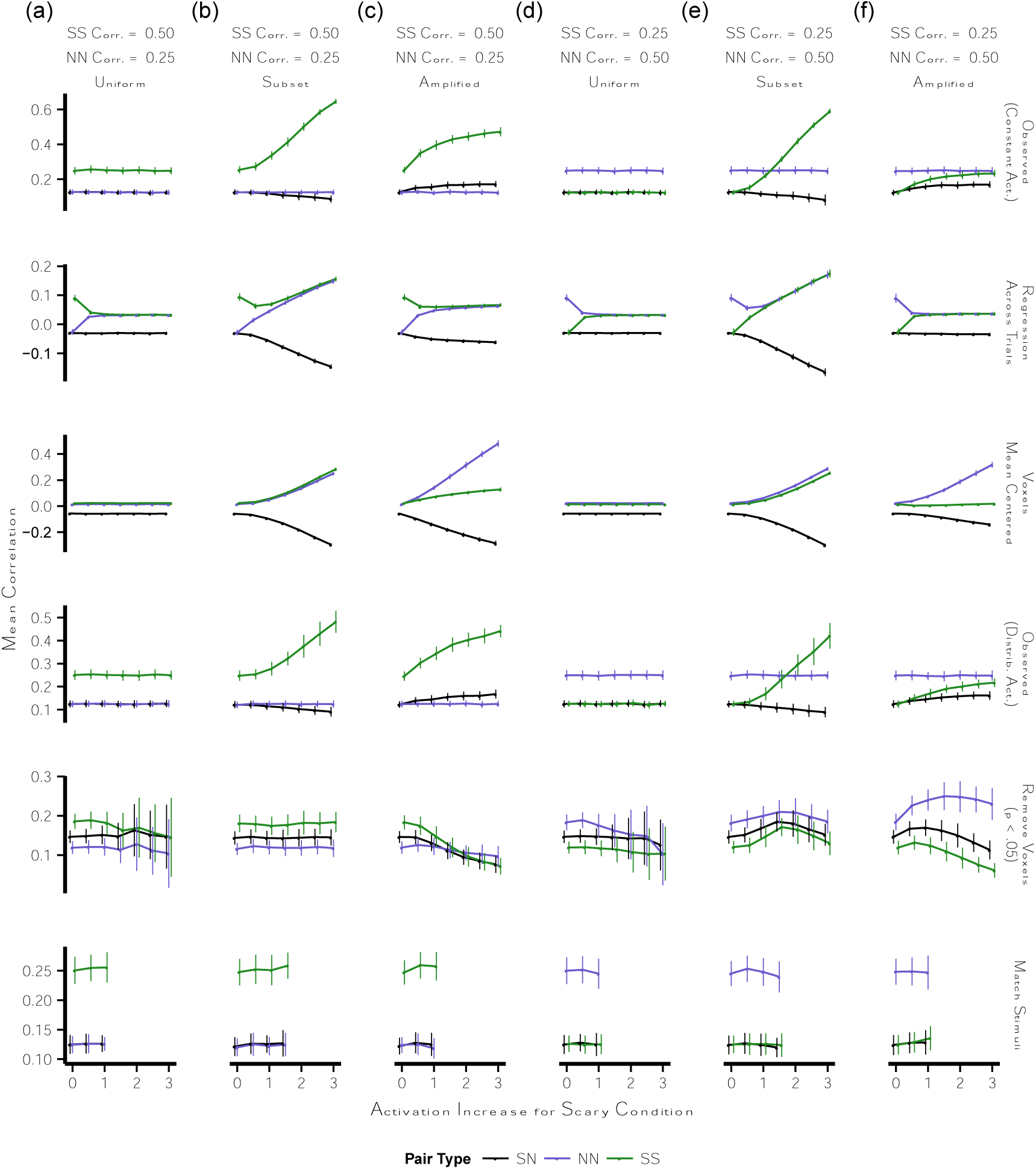
Observed and adjusted multivariate pattern similarity when condition produces simultaneous effects on activation and correlation. Points indicate the mean observed or adjusted correlation for pairs of stimuli at parametrically varying levels of an increase in activation for stimuli in the scary condition; note that 0 on the x-axis indicates no increase in activation. Error bars are standard deviation across 100 simulations. Activation was added to stimuli in the scary condition as **(a, d)** a uniform increase across all voxels (*uniform* case), **(b, e)** a uniform increase for 50% of the voxels (*subset* case), or **(c, f)** a stimulus-specific increase across all voxels equal to a multiple of the baseline response to each stimulus (*amplified* case). The underlying correlation across voxels was set to .25 for SN pairs and either **(a, b, c)** .50 for SS pairs and .25 for NN pairs or **(d, e, f)** .50 for NN pairs and .25 for SS pairs. ***Top-bottom***: (first row) observed correlations when the increase in activation was constant across stimuli; (second row) residuals after regressing correlation on activation across all pairs of stimuli; (thirid row) correlations between residuals after mean-centering each voxel; (fourth row) observed correlations when the increase in activation was distributed across stimuli according to a χ^2^ distribution with mean shown on the x-axis; (fifth row) observed correlations after removing voxels showing a significant difference in activation between scary and non-scary stimuli at p < .05; (sixth row) observed correlations after retaining only a subset of stimuli that are matched on mean activation across voxels. Panels ii and iii reflect adjustments relative to the observed data in panel i; panels v and vi reflect adjustments relative to the observed data in panel vi. *NN* = pairs of stimuli both drawn from the non-scary condition, *SS* = pairs of stimuli both drawn from the scary condition, *SN* = pairs of stimuli in which one was drawn from the scary condition and one was drawn from the non-scary condition, *corr.* = correlation, *act.* = activation.

**Figure S6.**
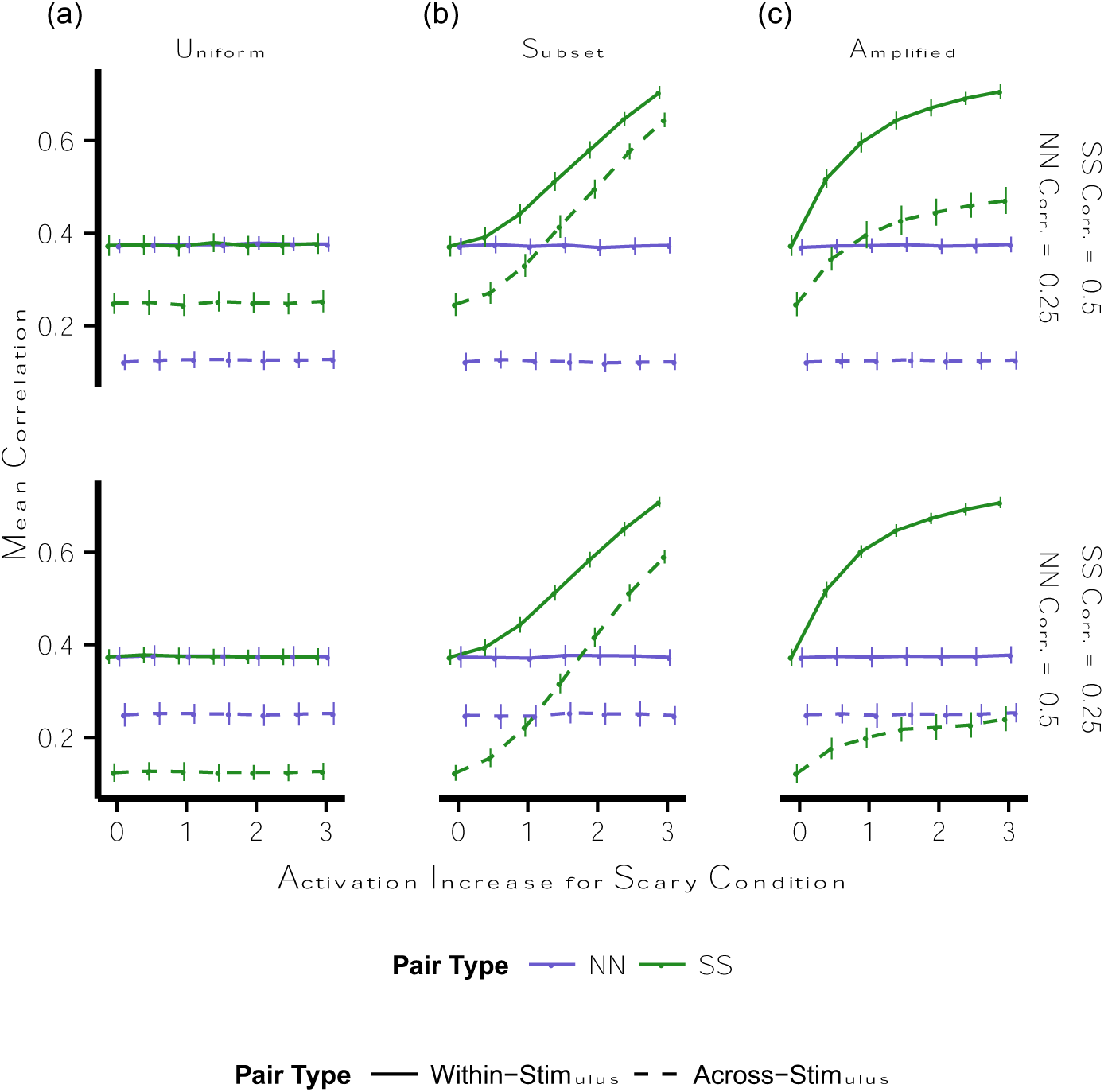
Observed and adjusted multivariate pattern similarity in a design including experimental and control cells when condition produces effects on activation in all cells and simultaneous effects on correlation in experimental cells. Points indicate mean adjusted correlation estimate for pairs of stimuli at parametrically varying levels of an increase in activation for stimuli in the scary condition; note that 0 on the x-axis indicates no increase in activation. Error bars are standard deviation across 100 simulations. Activation was added to stimuli in the scary condition as **(a)** a uniform increase across all voxels (*uniform* case), **(b)** a uniform increase for 50% of the voxels (*subset* case), or **(c)** a stimulus-specific increase across all voxels equal to a multiple of the baseline response to each stimulus (*amplified* case). ***Top-bottom***: the underlying correlation across voxels was set to .75 for within-stimulus pairs and either (top) .50 for across-stimulus SS pairs and .25 for across-stimulus NN pairs or (bottom) .50 for across-stimulus NN pairs and .25 for across-stimulus SS pairs. *NN* = pairs of stimuli both drawn from the non-scary condition, *SS* = pairs of stimuli both drawn from the scary condition, *corr.* = correlation.

## Notes

### Competing Interest Statement

The authors have declared no competing interest.

### Summary of Updates

Corrected verb-tense agreement in abstract

